# A global view of the RNA-binding and regulatory protein landscape in *Caenorhabditis elegans*

**DOI:** 10.1101/2024.11.07.622524

**Authors:** John D. Laver, Maida Duncan, John A. Calarco

## Abstract

Post-transcriptional regulation of gene expression is essential for the correct development and functioning of an organism. This regulation is coordinated by a collection of proteins that work together to determine an RNA’s post-transcriptional fate. Here, we provide a global overview of the RNA regulatory protein landscape in *Caenorhabditis elegans*, to provide insight into the coordination of post-transcriptional regulatory activities in the context of a multicellular organism. First, we have curated a comprehensive list of all known and putative RNA regulatory proteins encoded in the *C. elegans* genome, classified based on domain and functional annotations and published experimental data. Second, using protein-protein interaction data in the STRING database, we created a putative RNA regulatory protein interaction network that highlighted known RNA regulatory complexes, and leveraged this network to identify an additional 138 known and putative RNA regulators previously unannotated in *C. elegans*. Finally, we examined the tissue- and developmental-stage-specific expression of RNA regulators using published transcript expression data, which revealed strong expression in the gonad for a majority, as well as dozens expressed specifically in each of the major somatic *C. elegans* tissues. Taken together, this work will provide a valuable resource for future studies of RNA biology in *C. elegans*.

## Introduction

Precise regulation of gene expression is essential for nearly all cellular processes. Cells regulate gene expression before, during, and after transcription and translation by a diversity of mechanisms. At the post-transcriptional stage of gene regulation, a variety of proteins have roles in mediating RNA regulatory processes such as alternative splicing, polyadenylation, RNA editing, RNA transport, RNA localization, translation, and RNA degradation (Nishikura 2016; Roundtree et al. 2017; Bovaird et al. 2018; Heck and Wilusz 2018; Williams et al. 2018; Karousis and Mühlemann 2019; Ule and Blencowe 2019; Das et al. 2021; Machado de Amorim and Chakrabarti 2022; Boreikaitė and Passmore 2023; Bourke et al. 2023). For example, RNA-binding proteins (RBPs) interact directly with RNA via well-characterized RNA-binding domains (RBDs), forming ribonucleoprotein complexes (RNPs) that are responsible for many aspects of RNA metabolism (Corley et al. 2020).

In addition to these canonical RBPs, other proteins also contribute to post-transcriptional regulation, either by working together with RBPs via protein-protein interactions, as for example in molecular machines such as the ribosome and spliceosome, or by directly regulating or modifying the RBPs themselves, for example via post-translational modifications (Baßler and Hurt 2019; Hofweber and Dormann 2019; Wilkinson et al. 2020; Velázquez-Cruz et al. 2021; Ripin and Parker 2023). Finally, in recent years experimental approaches such as RNA interactome capture have identified proteins presumed to directly interact with RNA through less conventional, and still poorly characterized, binding modalities (Baltz et al. 2012; Castello et al. 2012; Castello et al. 2016; Perez-Perri et al. 2021; Ray et al. 2023). These non-canonical RBPs have greatly expanded the landscape of putative post-transcriptional regulatory factors (Hentze et al. 2018).

Due to the extensive involvement of RNA regulatory factors in controlling gene expression, their altered function has been implicated in many diseases. In several cancer types, for example, dysregulation of RBPs is associated with changes in expression of tumor suppressor genes and oncogenes (Pereira et al. 2017; Zhao et al. 2022). The dysregulation of RBPs is also often associated with neurodegenerative diseases (Cookson 2017; Wolozin and Ivanov 2019; Prashad and Gopal 2021). For example, Spinal Muscular Atrophy (SMA), a disease characterized by the gradual death of motor neurons, is caused by the loss of the RBP splicing factor Survival of Motor Neuron 1 (SMN1) (Mercuri et al. 2022). Other neurodegenerative diseases associated with the dysregulation of RBPs include Amyotrophic Lateral Sclerosis (ALS) and fragile X-associated tremor/ataxia syndrome (FXTAS) (Hagerman and Hagerman 2015; Hardiman et al. 2017). The role of RBP dysfunction in these diseases highlights the need for a thorough understanding of how these proteins operate together within the cell, as well as how their activities are coordinated across tissues in complex multicellular organisms.

To gain a complete picture of RNA regulatory protein function in an organism, an essential starting point is to have: (i) a complete list of all the known and putative RNA regulatory factors encoded in its genome; (ii) an understanding of how these factors cooperate with one another; and (iii) knowledge of where and when in the organism they are acting. The nematode, *Caenorhabditis elegans*, is a complex multicellular animal with a diversity of tissues and cell types, which serves as a powerful model system for studies of gene regulation *in vivo*.

Here, we aim to provide a global overview of the RNA regulatory protein landscape in *C. elegans*. First, using a combination of computational approaches, we compiled an exhaustive list of 2,152 known and putative *C. elegans* RNA regulatory factors, including canonical RBPs, their associated factors and regulators, and novel non-canonical RBPs identified by RNA interactome capture. Second, we used reported protein-protein interaction data to investigate how these factors might cooperate in their regulatory activities, and to identify an additional 138 putative RNA regulators not currently annotated with RNA-related functions in *C. elegans*. Finally, we analyzed published gene expression data from different developmental stages and tissue types to reveal how these proteins might contribute to *C. elegans* tissue development and function, which pointed to an especially important role for post-transcriptional regulation of gene expression in the germline. Together, this work will serve as an important resource for further studies of post-transcriptional regulation of gene expression in *C. elegans*, as well as a framework for similar studies in other organisms.

## Materials and Methods

### Curation of RNA-binding and regulatory proteins

RNA-binding proteins and other proteins with roles in RNA regulation and function were defined based on: the presence of canonical sequence- or structure-specific RNA-binding domains (Class 1); the presence of other non-specific RNA-associating or RNA-metabolism-related domains (Class 2); annotation with RNA metabolism-related Gene Ontology (GO) terms or UniProt keywords, or identification in previous *C. elegans* RBP curation studies (Tamburino et al. 2013) (Class 3); or identification as part of RNA interactome capture studies (Matia-González et al. 2015; Esmaillie et al. 2019) (Class 4).

Lists of sequence- or structure-specific RNA-binding domains, and non-specific RNA-binding and RNA-metabolism-related domains, were compiled by their InterPro database accession (https://www.ebi.ac.uk/interpro/), based on existing literature (Cook et al. 2011; Gerstberger et al. 2014; Loedige et al. 2014). RNA regulation-related GO terms and UniProt keywords were compiled by searching for terms (via the QuickGO web interface: https://www.ebi.ac.uk/QuickGO/) or keywords (via the UniProt website: https://www.uniprot.org/keywords/) containing “RNA” or words describing co- and post-transcriptional processes, excluding terms related to RNA anabolic processes and transcription. The annotated domains, GO terms and UniProt keywords used are presented in Table S1.

Complete lists of *C. elegans* genes annotated with these domains, GO terms (including child terms) or UniProt keywords were compiled based on the annotations assigned in the *C. elegans* proteome entry on the UniProt database (UniProt Proteome ID: UP000001940; proteome and relevant annotations downloaded September 5, 2023).

Proteins that were identified as RNA-binding proteins by a previously published compilation of *C. elegans* RNA-binding proteins (Tamburino et al. 2013), but which were not present in our lists of genes annotated with relevant domains, GO terms, and UniProt keywords, were manually reviewed for evidence of RNA regulatory function, and ten such proteins were added to our list.

A list of *C. elegans* proteins identified as RNA-associated by RNA interactome capture experiments was compiled by taking the union of all such proteins identified in the two published studies performed to date in *C. elegans* (Matia-González et al. 2015; Esmaillie et al. 2019). In Esmaillie *et al*. 2019 only proteins that were identified experimentally as RNA-binding in wild-type *C. elegans* were included in our protein list.

Our complete list of 2,152 RNA-binding and regulatory proteins is presented in Table S2, including domain and functional annotation information, the presence or absence in RNA interactome capture studies, and protein Class designations.

### Protein-protein interaction network: generation, clustering, GO enrichment analysis, and network expansion

The RNA regulatory protein-protein interaction network was generated with Cytoscape software v3.10.1 (Shannon et al. 2003) using interaction data in the STRING database (https://string-db.org/) (Szklarczyk et al. 2019; Szklarczyk et al. 2023), via the Cytoscape stringApp v2.0.1 (Doncheva et al. 2019; Doncheva et al. 2023). Software and database versions used were current as of September 10, 2023. Specifically, the entire list of 2,152 RNA regulatory proteins (all of Classes 1, 2, 3 and 4) was entered as a STRING protein query in the stringApp, using the corresponding WormBase ID gene identifiers (https://wormbase.org) (Sternberg et al. 2024). The stringApp query settings were for network type – physical subnetwork; confidence (score) cutoff – 0.90 (termed “highest confidence” by STRING); and maximum additional interactors – 0. This generated a network for 2,139 proteins with recognized IDs that were present in the STRING database, 944 of which had 7,688 reported interactions with other RNA regulatory proteins (Figure 2, Figure S1, File S1, File S2).

Clustering was performed on the protein-protein interaction network to identify highly interconnected groups of proteins, using the Cytoscape app clusterMaker2 (Morris et al. 2011; Utriainen and Morris 2023). Specifically, all 944 RNA regulatory proteins with interactions in the STRING network were subjected to clustering, using the clusterMaker2 MCL clustering algorithm, with the “granularity” parameter set to 2.0, “array source” set as stringdb::score, and remaining settings left as default.

GO enrichment analysis was performed on each of the protein-protein interaction network clusters that consisted of at least 10 proteins (22 clusters total). This analysis was carried out using the g:Profiler web interface (version e110_eg57_p18_4b54a898) (Kolberg et al. 2023). Specifically, WormBase ID gene identifiers were input for each of the 22 clusters being analyzed, as a multiquery, with the organism database selected as *Caenorhabditis elegans* (PRJNA13758) from WormBase ParaSite. Enrichments were assessed against the background of all 2,139 genes that were recognized in our STRING protein-protein interaction network query, using the g:SCS multiple testing correction method and significance threshold of 0.05. The full results of the GO enrichment analysis are presented in Table S3. For selected terms, fold enrichment values were calculated manually for each GO term/cluster, as [(number of genes in cluster annotated with GO term)/(number of genes in cluster)]/[(number of genes in background annotated with GO term)/(number of genes in background)].

To identify additional proteins with potential roles in RNA regulation, the RNA regulatory protein-protein interaction network was expanded to include additional proteins, based on interactions with RNA regulatory proteins in Classes 1, 2 or 3. Specifically, in the network of 2,139 proteins, Class 1, 2 and 3 proteins were selected, and additional interacting proteins were identified using the “Expand network” tool of the stringApp (Doncheva et al. 2019; Doncheva et al. 2023), with the “selectivity” setting at 0.5 and the “number of new nodes” to be added set at 1,000. This resulted in the addition of 200 proteins to the network. After creating a new protein-protein interaction network with just Class 1, 2 and 3 proteins and these 200 new proteins, the 200 new proteins were filtered to only include those with at least two interactions in the new network (i.e. at least one interaction with a Class 1, 2, or 3 protein, and at least one interaction with either another Class 1-3 protein or another new protein in the set of 200), resulting in 138 new proteins, which we called Class 5 proteins, added to the original set of 2,152 RNA regulatory proteins in Classes 1-4 (Figure 3, Figure S2, Table S4, File S3, File S4). To examine the functions of the proteins present in Class 5, we performed GO enrichment analysis using the g:Profiler web interface (version e111_eg58_p18_b51d8f08) (Kolberg et al. 2023). Specifically, WormBase ID gene identifiers were input as a single query, with the organism database selected as *Caenorhabditis elegans* (PRJNA13758) from WormBase ParaSite. Enrichments were assessed against the background of all annotated genes in the *C. elegans* genome, using the false discovery rate multiple testing correction method and significance threshold of 0.05. The full results of the GO enrichment analysis are presented in Table S5. For selected GO terms, fold enrichment values were calculated manually as [(number of genes in query annotated with GO term)/(number of genes in query)]/[(number of genes in genome annotated with GO term)/(number of genes in background)].

### Analysis of RNA regulatory gene expression: datasets used, clustering, analysis of evolutionary conservation, and GO enrichment analysis

Data describing transcript expression for RNA regulatory proteins in Classes 1, 2, and 3 were extracted from published datasets describing tissue-specific gene expression during embryogenesis (Warner et al. 2019), and at L2 larval (Cao et al. 2017), L4 larval (Gracida et al. 2017), and adult (Kaletsky et al. 2018) stages of development. For the L2 larval expression data, TPMs for the tissues examined were extracted from tissue-level consensus expression profiles presented in Table S3 of Cao et al., 2017, and TPMs for gonad cell types examined were extracted from the cell-type consensus expression profiles presented in Table S4 of Cao et al., 2017. For the embryogenesis data, we focused on the results of the fuzzy k-means clustering presented in Warner et al., 2019, which assigned genes to clusters based on their primary tissue or temporal pattern of expression; cluster assignments were extracted from Table S5 of Warner et al., 2019 (“BestCluster”), and descriptions of the tissue or temporal expression patterns of different clusters were derived from the labels in the t-SNE plot in Figure 2 of Warner et al., 2019. For L4 larval expression data, genes enriched in neurons, intestine or muscle were extracted from the lists of genes presented in Table S1 of Gracida et al., 2017, as neuron-enriched, intestine-enriched, or muscle-enriched. For adult expression data, genes enriched in neurons, intestine, muscle, or hypodermis were extracted from the lists of tissue-enriched genes presented in Table S8 of Kaletsky et al., 2018, excluding any genes that were described in Table S8 as enriched in more than one tissue.

RNA regulatory proteins were clustered based on the tissue expression profiles (gonad, intestine, neurons, glia, pharynx, muscle, hypodermis) of their encoding transcripts at the L2 larval stage, from Cao et al., 2017. Specifically, after excluding genes that were not expressed with TPM ≥ 0.5 in at least one tissue, expression values for each gene in all tissues were normalized to the tissue with the highest expression. These data were used to cluster all genes into 13 clusters by k-means clustering, using the ‘kmeans’ function in the ‘stats’ package in R, setting the maximum number of iterations (‘iter.max’) to 20 and ‘nstart’ to 1000.

For analysis of the evolutionary conservation of *C. elegans* RNA regulatory proteins, information about homologous proteins in other model organisms from the Alliance of Genome Resources (https://www.alliancegenome.org/) was downloaded via the AllianceMine web interface (https://www.alliancegenome.org/bluegenes/alliancemine) (Smith et al. 2012; Alliance of Genome Resources Consortium 2024). For our analysis, *C. elegans* proteins were considered to have a direct ortholog in another model organism if the second species had a homolog to the *C. elegans* protein that was defined by the Alliance of Genome Resources analysis as being the best predicted ortholog in that species, with the *C. elegans* protein also being the best predicted ortholog among *C. elegans* proteins for that particular homolog.

GO enrichment analysis was performed on each of the 13 L2 larval expression clusters. This analysis was carried out using the g:Profiler web interface (version e110_eg57_p18_4b54a898) (Kolberg et al. 2023). Specifically, WormBase ID gene identifiers were input for each of the 13 clusters being analyzed, as a multiquery, with the organism database selected as *Caenorhabditis elegans* (PRJNA13758) from WormBase ParaSite. Enrichments were assessed against the background of all 1,353 Class 1-3 genes, using the false discovery rate multiple testing correction method and significance threshold of 0.05. To simplify the results, GO terms with fewer than 10 annotated genes in the background set were excluded. The full results of the GO enrichment analysis are presented in Table S6. For selected terms, fold enrichment values were calculated manually for each GO term/cluster, as [(number of genes in cluster annotated with GO term)/(number of genes in cluster)]/[(number of genes in background annotated with GO term)/(number of genes in background)].

For analysis of the expression of genes with functions in RNA splicing, shown in Figure 6B, the list of all *C. elegans* genes annotated with the GO term “regulation of RNA splicing” (GO:0043484) was obtained from WormBase [downloaded February 12, 2024]. Expression of these genes at the L2 larval stage, from Cao et al., 2017, was analyzed in the six non-gonadal tissues by normalizing expression values for each gene in all non-gonadal tissues to the non-gonadal tissue with the highest expression.

For analysis of expression in gonad cell types at the L2 larval stage from Cao et al., 2017, shown in Figure 5, expression values for each gene in all gonad cell types were normalized to the gonad cell type with the highest expression. For more detailed analysis of the expression of germline-enriched RNA regulatory genes in non-gonadal tissues, L2 expression cluster 7 – which among all clusters and tissues/cell-types was most highly expressed in germline – was examined. Specifically, after excluding genes that were not expressed with TPM ≥ 0.5 in at least one non-gonadal tissue, expression values for each gene in the non-gonadal tissues were normalized to the non-gonadal tissue with the highest expression. These normalized expression data for the six non-gonadal tissues (intestine, neurons, glia, pharynx, muscle, hypodermis) were used to further cluster the genes in L2 expression cluster 7 into 6 sub-clusters by k-means clustering, using the ‘kmeans’ function in the ‘stats’ package in R, setting the maximum number of iterations (‘iter.max’) to 20 and ‘nstart’ to 1000.

All heatmaps displaying expression and tissue enrichment data were generated in R using the ComplexHeatmap package (Gu et al. 2016; Gu 2022).

### Software for statistical tests

Fisher’s exact tests and Wilcoxon rank-sum tests were performed using the ‘fisher.test’ and ‘wilcox.test’ functions, respectively, in the R ‘stats’ package. Two-sample Anderson-Darling tests for comparing distributions were performed using the ‘ad_test’ function in the R package ‘twosamples’ v2.0.1 (Dowd 2023).

## Results and Discussion

### A curated set of RNA-binding and regulatory proteins in C. elegans

We set out to compile a comprehensive and up-to-date collection of *C. elegans* proteins with known or putative roles in co-transcriptional or post-transcriptional RNA metabolism and function, building on previous work by Tamburino et al. (2013), and others (Cook et al. 2011). Proteins may participate in the regulation of RNA metabolism and function through a variety of mechanisms, including directly binding RNA via specific sequence- or structure-based *cis*-elements, directly binding to RNA in a non-sequence-specific manner, associating indirectly with RNA via protein-protein interactions with RNA-binding proteins or as part of large macromolecular RNA regulatory machines (such as the ribosome and spliceosome), or by mediating post-translational modifications that regulate the activity of RNA-associated factors.

We used a variety of resources to systematically identify all *C. elegans* proteins with known or predicted functions in co- and post-transcriptional RNA regulation and function (Figure 1A and Table S1). First, we compiled all *C. elegans* proteins (427 in total) containing annotated domains with established roles in specific or non-specific RNA binding or RNA metabolism (Cook et al. 2011; Gerstberger et al. 2014), using annotations from the InterPro database. Second, we used the functional annotations defined in the Gene Ontology (GO) database and UniProt to compile all proteins (1,292 in total) annotated as having roles in RNA regulation; notably, we excluded GO terms related to RNA anabolic processes, as this would have resulted in the inclusion of a large number of transcriptional regulators, which are not the focus of this analysis. The RNA-interacting domains, GO terms and UniProt keywords used to compile these lists are summarized in Figure 1A, with the full details in Table S1. We also carefully examined the previously published list of *C. elegans* RNA-binding proteins (Tamburino et al. 2013) for any proteins that were not yet identified by our searches but had strong evidence of a function in RNA regulation, resulting in the addition of 10 further proteins. Comparison of the proteins collected from these different sources (Figure 1B) revealed, as expected, that the majority of proteins containing RNA-binding or RNA-metabolism-related domains were also functionally annotated in the GO or UniProt databases as having roles in RNA regulation (376 of 427, 88% of domain-annotated proteins). There were an additional 926 proteins identified that were annotated with RNA regulatory functions by the GO database, UniProt, or previous reports (Tamburino et al. 2013) that did not contain RNA binding or regulatory domains.

**Figure 1.**
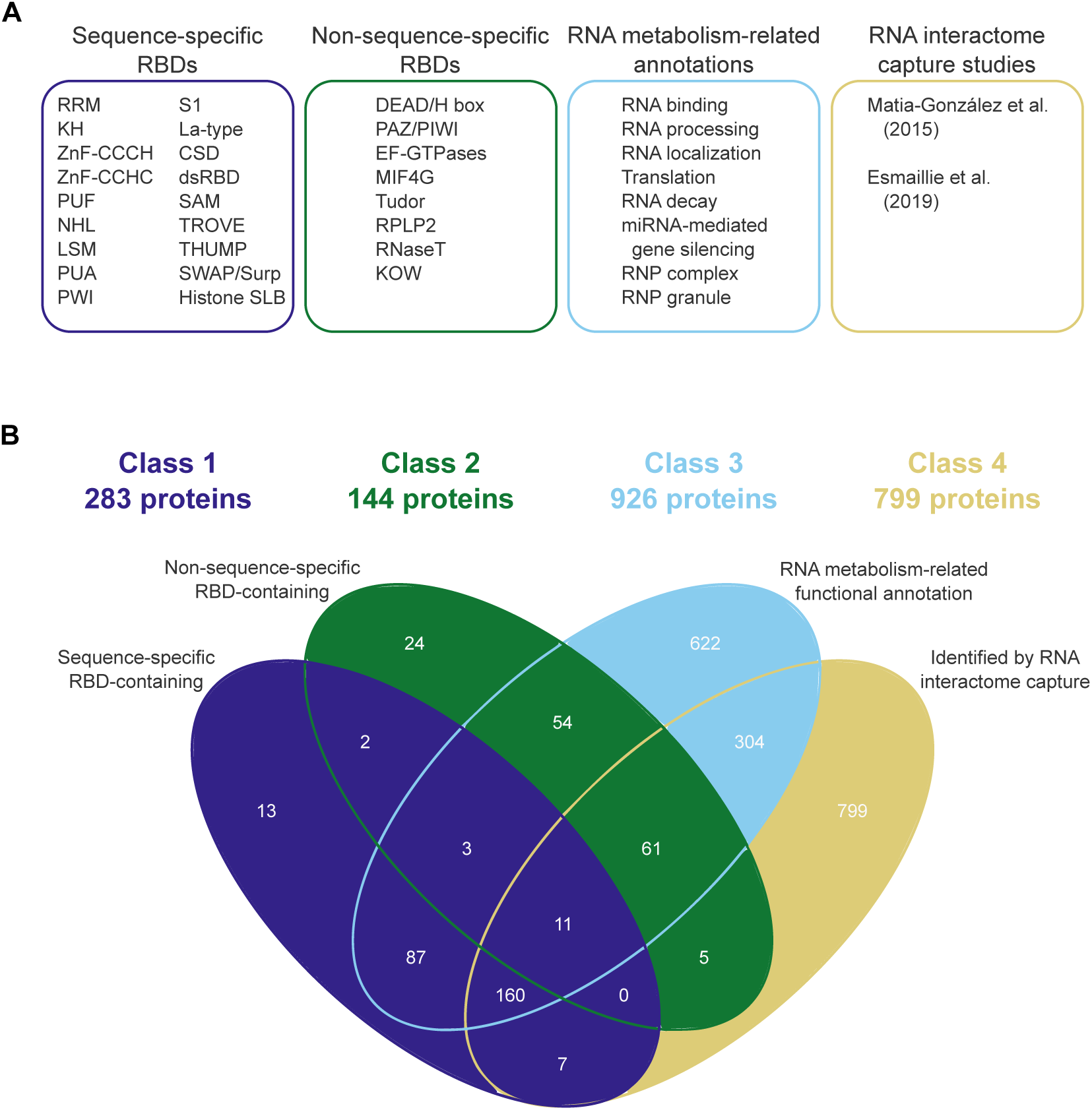
Compiling a comprehensive list of RNA regulatory proteins in *C. elegans*. **(A)**Summary of (i) sequence- or structure-specific RNA-binding domains (RBDs), (ii) non-specific RBDs or RNA metabolism-related domains, (iii) RNA metabolism-related functional annotations, and (iv) RNA interactome capture studies, that were used to compile a comprehensive list of *C. elegans* RNA regulatory proteins; full details are presented in Table S1. **(B)** Comparison of protein lists derived from categories in (A), and categorization into four non-overlapping groups comprising 2,152 total proteins: Class 1 (sequence- or structure-specific RNA-binding proteins), Class 2 (non-specific RNA-binding proteins), Class 3 (non-RNA binding proteins with roles in RNA regulation), and Class 4 (putative RNA-binding proteins identified by RNA interactome capture).

To complement this set of *bona fide* RNA regulatory proteins, we also compiled a set of 1,347 proteins that have been experimentally identified to be associated with RNA in *C. elegans* by RNA interactome capture methods (Matia-González et al. 2015; Esmaillie et al. 2019). These experiments used UV crosslinking to covalently link RNA to bound proteins, followed by isolation of bulk polyadenylated RNA and mass spectrometry-based identification of associated proteins. Of the proteins identified by these studies, 548 (41%) were already identified through the domain or functional annotation searches described above, whereas the remaining 799 (59%) were only identified by RNA interactome capture (Figure 1B). This latter group of proteins represent potentially under-explored candidates to connect to known RNA metabolic pathways via other lines of evidence.

Combining the proteins collected from all these sources resulted in a list of 2,152 unique *C. elegans* RNA regulatory factors; the complete list along with information regarding relevant annotations is presented in Table S2. We categorized these proteins to reflect the various modes of RNA interaction described above, by subdividing them into four mutually exclusive, hierarchically-organized classes (Figure 1A and 1B; Table S2): Class 1 (283 proteins) were defined as those containing canonical sequence- or structure-specific RNA-binding domains, such as the RRM, KH domain, ZnF domains, and dsRBD; Class 2 (144 proteins) as those containing non-specific RNA-binding or metabolism-related domains (excluding proteins already part of Class 1), such as the DEAD/H box and PAZ/PIWI domains; Class 3 (926 proteins) as those annotated with RNA-related functions (excluding proteins already part of Classes 1 or 2), which includes many components of large RNA metabolic complexes such as the ribosome, as well as some post-translational modifiers; and Class 4 (799 proteins) as those identified only as part of RNA interactome capture studies (i.e. excluding proteins already part of Classes 1, 2, or 3). Taken together, this list provides the most comprehensive atlas of proteins to date that are associated with various aspects of RNA regulation and function in *C. elegans*.

### A putative protein-protein interaction network reveals connectivity within and between known RNA regulatory complexes

To gain insight into the connections between RNA regulatory proteins and processes, we next used protein interaction data curated in the STRING database (Szklarczyk et al. 2019) to create a putative protein-protein interaction (PPI) network using all 2,152 RNA regulatory factors identified above, which we visualized using Cytoscape (Shannon et al. 2003) (Figure 2A, Figure S1, File S1, and File S2; see Materials and Methods for details). Of the total set of 2,152 RNA regulatory proteins, 2,139 were found in the STRING database, of which 944 were reported to interact with other proteins within the set of RNA regulators, under the stringent confidence criteria we selected. Of these, 664 proteins formed a single interconnected network comprising 7,247 interactions.

**Figure 2.**
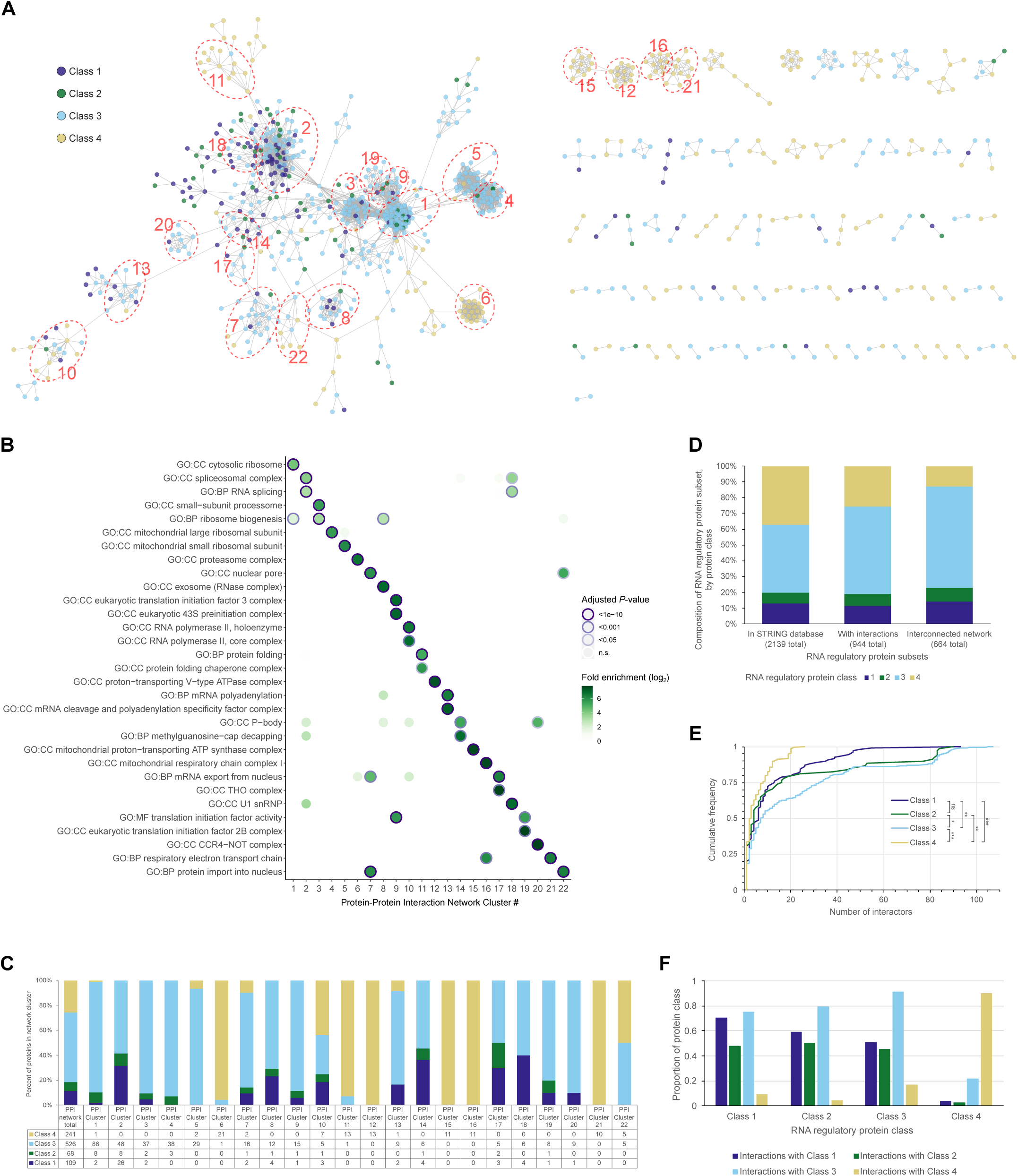
Global overview of reported protein-protein interactions among *C. elegans* RNA regulatory proteins. **(A)** Network diagram of reported protein-protein interactions among all classes of RNA regulatory proteins, with nodes representing proteins, colour-coded by RNA regulatory protein class, and edges representing protein-protein interactions reported in the STRING database. 944 of 2,152 total RNA regulatory proteins have reported interactions with other RNA regulators and are shown here; 664 proteins form a fully interconnected network (left side). Clustering was performed to identify groups of highly interacting proteins; 22 clusters with 10 or more proteins are delineated by the numbered dashed circles. Network image with nodes labelled with corresponding gene names is presented in Figure S1. **(B)** Summary of GO term enrichment analysis performed on each of the protein-protein interaction clusters, against the background of the entire list of 2,152 RNA regulatory proteins. Fold enrichment and adjusted *P*-value for each GO term/cluster pairing are indicated by the fill and outline of the circles, respectively. In general each cluster corresponds to a specific known protein complex. Full results of the GO enrichment analysis are presented in Table S3. **(C)** Composition of protein-protein interaction clusters by regulatory protein class; proportions of each class comprising the 944 proteins in the overall network are included for comparison. **(D)** Percent composition, by RNA regulatory protein class, of (i) the entire set of RNA regulatory proteins found in the STRING database (2,139 total), (ii) the subset of RNA regulatory proteins reported as having interactions with other RNA regulators (all proteins shown in network in panel (A); 944 total), and (iii) the subset of RNA regulatory proteins that form a single large fully interconnected protein-protein interaction network (left side of panel (A); 664 total). **(E)** Cumulative distributions of the number of interactors per protein for each of the RNA regulatory protein classes. Statistical significance between distributions was calculated by two-sample Anderson-Darling test and is indicated: \**P* < 0.05; \*\**P* < 0.01; \*\*\**P* < 0.001; ns – not significant (*P* > 0.05). **(F)** For all the proteins in each RNA regulatory protein class that have reported interactions, the proportion that interact with at least one protein from each of the four classes is shown.

To formally identify complexes and other groups of cooperating proteins that were highly interconnected via protein-protein interactions, we performed network clustering analysis (see Materials and Methods for details). Within the subset of 944 interacting proteins, this identified 517 proteins that grouped into 22 clusters containing ten or more proteins (PPI clusters 1-22), and an additional 158 proteins that were part of 23 clusters with between five and nine proteins (PPI clusters 23-45; Figure 2, Table S2).

We systematically identified functions and known complexes associated with the 22 clusters of at least ten proteins by performing GO term enrichment analysis against the background of the total list of 2,139 RNA regulatory proteins found in the STRING database. This revealed that the different clusters largely corresponded to specific known protein complexes (Figure 2B, Table S3). The largest clusters, for example, corresponded to the cytosolic ribosome and associated factors (PPI cluster 1; 97 proteins), the spliceosome (PPI cluster 2; 82 proteins), ribosome biogenesis machinery (PPI cluster 3; 41 proteins), and mitochondrial ribosome large and small subunits (PPI clusters 4 and 5; 41 and 31 proteins, respectively). Additional clusters corresponded to components of RNA polymerase II that possess RNA-binding domains or have secondary roles in post-transcriptional regulation or non-coding RNA function (PPI cluster 10; 16 proteins), the U1 snRNP (PPI cluster 18; 10 proteins), the cleavage and polyadenylation specificity factor complex (PPI cluster 13; 12 proteins), the nuclear pore complex (PPI cluster 7; 21 proteins), nucleocytoplasmic transport complexes (PPI clusters 17 and 22; 10 proteins each), translation initiation factor complexes (PPI clusters 9 and 19; 17 and 10 proteins, respectively), the exosome (PPI cluster 8; 17 proteins), mRNA decapping machinery (PPI cluster 14; 11 proteins), and the CCR4-NOT complex (PPI cluster 20; 10 proteins).

Most of these RNA regulatory complex-related clusters contained proteins from all of Classes 1, 2 and 3, in varying proportions (Figure 2C), although some clusters were almost entirely composed of Class 3 proteins (e.g. the ribosome-related clusters). There was a notable absence of Class 4 proteins in these RNA regulatory complex-related clusters, with the exception of PPI clusters 10 (RNA polymerase II) and 22 (protein import into the nucleus). In contrast, the remaining clusters of ten or more proteins not listed above were composed almost entirely of Class 4 proteins (Figure 2C) and corresponded to complexes with no direct known role in RNA regulation or function, consistent with the novelty of the RNA-binding activity of the proteins in this class. These Class 4 protein-containing clusters included protein chaperones (PPI cluster 11; 14 proteins), the proteasome (PPI cluster 6; 22 proteins), V-type ATPase (PPI cluster 12; 13 proteins), and mitochondrial respiratory chain complexes (PPI clusters 15, 16 and 21; 11, 11, and 10 proteins, respectively).

Collectively, this functionally annotated protein-protein interaction network formed by our curated RNA regulatory proteins allows for a survey of the interactions among these factors to rapidly query potential connections across different layers of RNA metabolism, while the subset of RBPs and RNA regulatory factors that lack connections to the broader network may represent less well-characterized proteins to study in more detail.

### Proteins identified by RNA interactome capture without previously characterized roles in RNA regulation have few interactions with other RNA regulatory proteins

Given the apparent partitioning between Class 1, 2, and 3 proteins and Class 4 proteins revealed by our clustering analysis, we more closely examined the presence and connectivity of each class of proteins within the network. We first assessed, compared to the total set of 2,139 proteins found in the STRING database, the proportions of each protein class in (i) the set of 944 proteins reported to interact with other RNA regulators (all proteins shown in Figure 2A), and (ii) the set of 664 proteins forming a single interconnected network (left-hand side of Figure 2A). This revealed that, among the set of 944 proteins, Class 3 proteins were significantly over-represented, whereas Class 4 proteins were significantly under-represented (Figure 2D) (Fisher’s Exact Test: Class 3 odds ratio = 2.59, *P*-value < 10^-25^; Class 4 odds ratio = 0.39, *P*-value < 10^-23^). This trend was also apparent, and even more striking, when we considered the proportions of each protein class among the set of 664 proteins forming a single interconnected network, and among this protein set, Class 2 proteins were also significantly over-represented (Figure 2D) (Fisher’s Exact Test: Class 2 odds ratio = 1.59, *P*-value < 0.01; Class 3 odds ratio = 3.58, *P*-value < 10^-39^; Class 4 odds ratio = 0.16, P-value < 10^-60^). Together this indicates that, compared to Class 1, 2, and 3 proteins, Class 4 proteins are less likely to have reported interactions with other proteins in the full set of 2,139 RNA regulators. In contrast, the over-representation of Class 3 proteins likely reflects the fact that this class was compiled, in part, by identifying proteins that were known to be part of multi-subunit RNA regulatory complexes.

We next focused on the set of 944 interacting proteins and compared the number of interactions for proteins in the different classes (Figure 2E). Overall, the number of reported interactions for any given protein ranged from 1 to 107. Class 3 proteins tended to have the highest number of interactions per protein (median = 7 and 75^th^ percentile = 32.75), followed by Class 1 and Class 2 proteins (Class 1 median = 6 and 75^th^ percentile = 13; Class 2 median = 4 and 75^th^ percentile = 16), and finally Class 4 proteins, which tended to have the fewest interactions (median = 3 and 75^th^ percentile = 8). We confirmed that the overall distributions of the number of interactions were significantly different for Class 3 compared to either Class 1 or 2 (two-sample Anderson-Darling test *P* < 0.05), which were similar, and that Class 1, 2, and 3 all had significantly different distributions than Class 4 (two-sample Anderson-Darling test *P* < 0.01) (Figure 2E). This trend is consistent with that described above of the proportions of the different protein classes present in the interacting sets of RNA regulatory proteins, and shows that even among those Class 4 proteins that are reported to interact with other RNA regulators, they tend to have fewer interactions compared to Classes 1, 2, and 3.

Finally, for the reported interactions, we examined whether there were any biases in the classes of the proteins that were interacting with each other. To do this, we considered all the proteins in each class with reported interactions and determined the proportion that interacted with proteins from each of the four classes (Figure 2F). Strikingly, Class 1, 2 and 3 proteins had many more reported interactions with other proteins in these three classes than with Class 4 proteins, and vice versa. The proportion of proteins in Classes 1, 2, and 3 that were reported to interact with Class 1 proteins ranged from 51%-71%, with Class 2 ranged from 46%-50%, and with Class 3 ranged from 75%-91%, whereas the proportion interacting with Class 4 was much lower ranging from 4%-17%. Conversely, Class 4 proteins were much more likely to have reported interactions with other Class 4 proteins (90% of Class 4) than with proteins from either Class 1, 2, or 3 (4%, 2%, and 22% of Class 4 proteins, respectively).

Together, these observations suggest that proteins identified as RNA regulators only by RNA interactome capture may not have canonical RNA regulatory functions comparable to those of proteins in Classes 1, 2, and 3. It is possible that some proportion of these Class 4 proteins engage with RNA as an artifact of the crosslinking approaches used, or bind RNA in a transient, spurious, or non-specific manner. For at least a subset of proteins identified by RNA interactome capture in other organisms, this conclusion is supported by recent follow-up studies that found either no RNA-binding activity for these proteins, or non-specific RNA-binding activity, when assayed *in vitro* (Vaishali et al. 2021; Ray et al. 2023). However, it has been reported that the RNA-binding activity of many proteins identified by RNA interactome capture is conserved in orthologs across multiple species (Matia-González et al. 2015; Hentze et al. 2018; Esmaillie et al. 2019), which might support a functional role for some of these protein-RNA interactions. One interesting possibility is that the RNA-binding activity of these proteins serves an inherently different role from the RNA regulators in Classes 1, 2 and 3. For example, there is recent evidence in other systems that, for some novel RNA-binding proteins identified by RNA interactome capture, the binding of an RNA ligand may function to regulate the protein’s activity (Horos et al. 2019; Huppertz et al. 2022). Additional studies of these novel Class 4 RNA-binding proteins in *C. elegans* will be needed to validate and better understand any function(s) of their RNA-binding activity.

### Expanding the RNA regulatory protein-protein interaction network identifies additional RNA regulators

To identify putative novel RNA regulators, as well as additional known factors that may have been missed by our initial curation efforts, we used STRING to expand our protein-protein interaction network to include proteins outside of the 2,152 RNA regulatory proteins we originally identified. STRING can be used to add proteins to the network, based on user-defined selectivity criteria that aim to find a balance between the number of interactions a new protein has with proteins in the existing network and the overall number of protein interactions of the new protein. We chose to expand our network in this manner based on protein-protein interactions with RNA regulatory proteins in Classes 1, 2 or 3, excluding Class 4, as the role of Class 4 proteins in RNA regulation remains uncertain, as discussed above. After using STRING to expand our network, we filtered the newly added proteins to include only those that had at least two interactions in the newly expanded network – at least one interaction with a Class 1, 2 or 3 protein, and at least one additional interaction, either with another Class 1-3 protein or with another newly added protein (see Materials and Methods for details). These steps resulted in the identification of 138 new proteins with potential roles in RNA regulation; we refer to these collectively as Class 5 proteins (Table S4). The newly expanded protein interaction network, comprising proteins in Classes 1, 2, 3 and 5, is shown in Figure 3A (also see Figure S2, File S3, and File S4).

**Figure 3.**
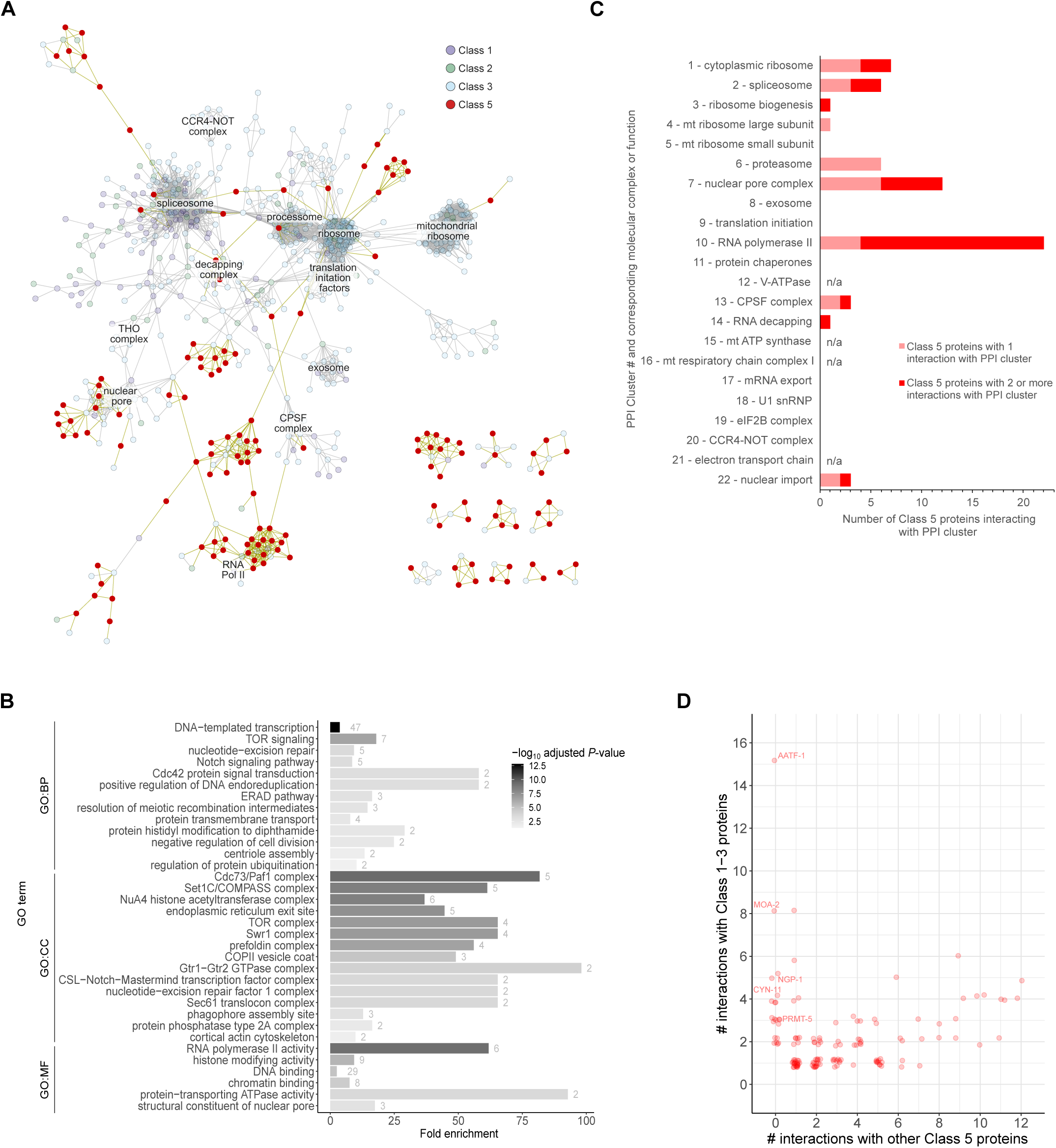
Identification of additional *C. elegans* RNA regulatory proteins by expansion of the RNA regulatory protein-protein interaction network. **(A)** Network diagram showing protein-protein interactions among RNA regulatory proteins in Classes 1, 2, and 3, as well as 138 additional putative RNA regulators that were identified on the basis of their reported interactions with Class 1-3 proteins. The 138 newly identified proteins (Class 5) are represented by red nodes, and their interactions are represented by dark yellow edges. The major molecular complexes identified by the clustering and GO enrichment analyses in Figure 2 are labelled. **(B)** Summary of GO term enrichment analysis performed on all Class 5 proteins, against the background of the entire *C. elegans* genome. Terms are organized based on the ontology to which they belong: biological process (GO:BP), cellular component (GO:CC), or molecular function (GO:MF). Full results of the GO enrichment analysis are presented in Table S5. **(C)** Number of Class 5 proteins interacting with each of the protein-protein interaction clusters reported in Figure 2. Class 5 proteins that interact with one protein from a given cluster are indicated by the lightly-shaded portion of the bars, and those that interact with two or more proteins in a given cluster are indicated by the darkly-shaded portion of the bars. Protein-protein interaction clusters are identified on the y-axis by the numbering presented in Figure 2, with the main molecular complex or function identified by GO enrichment analysis (Figure 2B) indicated. Clusters 12, 15, 16 and 21 were composed entirely of Class 4 proteins, and therefore were not expected to have any interactions with Class 5. **(D)** For each Class 5 protein, the number of interactions with Class 1-3 proteins is plotted against the number of interactions with other Class 5 proteins. Those proteins discussed in the main text are labelled. Note that points are slightly offset to allow better visualization of overlapping points.

To assess the known and potential roles of the Class 5 proteins and understand how they fit into the overall RNA regulatory landscape, we examined these proteins in two complementary ways. First, we performed GO enrichment analysis on the entire set of Class 5 proteins. This highlighted a variety of enriched complexes and pathways (Figure 3B, Table S5), which vary in the extent of their known connections to RNA regulatory processes.

In most cases, these complexes and pathways were already represented by at least one Class 1, 2 or 3 protein, but the newly added Class 5 proteins were not themselves annotated with any RNA regulatory functions. Nonetheless, several of these Class 5 protein complexes have well-known post-transcriptional regulatory roles. For example, via interactions with the Class 3 protein, TOR (LET-363), Class 5 proteins included several additional TOR complex components and regulators; TOR is known to impact multiple layers of post-transcriptional control through the regulation of post-translational modifications of other RNA regulatory proteins (Barbet et al. 1996; Brunn et al. 1997; Jefferies et al. 1997; Hu et al. 2015; Tang et al. 2018). Class 5 proteins also included 18 subunits of RNA polymerase II, largely due to their interaction with RNA polymerase II subunits RPB-4 (Class 3), RPB-7 (Class 1), and RPB-9 (Class 3), which have individual roles in post-transcriptional regulation (Choder 2004; Dahan and Choder 2013; Berkyurek et al. 2021; Richard et al. 2021); outside of its role in transcription, RNA polymerase II as a whole contributes to the regulation of a number of co-transcriptional RNA processing events (Moore and Proudfoot 2009). Additional examples of complexes or pathways that are enriched among Class 5 proteins and that have post-transcriptional functions include: the Sec61 translocon complex (two Class 5 proteins: SEC-61.B and SEC-61.G), which binds ribosomes and mediates the co-translational transport of newly synthesized polypeptides into the endoplasmic reticulum; diphthamide synthesis (two Class 5 proteins: DPH-1 and DPH-2), a conserved and essential post-translational modification to a histidine residue in eukaryotic translation elongation factor 2; and several subunits of the nuclear pore complex.

In addition to these complexes and pathways with post-transcriptional functions, there were a number of complexes enriched among Class 5 proteins that are known to mediate the function of non-coding RNAs in various aspects of DNA regulation (Smith and Shilatifard 2010; Khanduja et al. 2016; Bader et al. 2020; Statello et al. 2021). These included, for example, histone modifiers like the NuA4 histone acetyltransferase complex and Set1 methyltransferase complex, and DNA repair factors such as subunits of the nucleotide-excision repair complex 1.

Finally, there were some complexes enriched among Class 5 proteins whose roles in RNA regulation or function were not immediately obvious, as they were present due to interactions with Class 1, 2 or 3 proteins that have dual functions. For example, subunits of the COPII vesicle coat and ER exit site components – important in mediating the trafficking of cargo out of the ER – were identified as Class 5 proteins due to interactions with NPP-20, which is a component of the COPII coat but also functions in mRNA transport as a component of the nuclear pore. Similarly, Class 5 proteins included multiple subunits of the prefoldin complex – a protein chaperone that binds co-translationally to particular nascent polypeptide chains – as interactors of the prefoldin subunit PFD-3, which is annotated as polysome-associated and categorized as a Class 3 protein. Based on their most well-characterized functions neither of these complexes appear likely to have direct roles in RNA regulation. However, a survey of the literature reveals that both have in fact been shown to impinge on post-transcriptional control in other organisms: components of the ER exit site and COPII vesicle coat have been found to interact with and regulate the dynamics of stress granules and P-bodies (Aguilera-Gomez et al. 2017; Lee et al. 2020; Milano et al. 2024), whereas the prefoldin complex has been shown to impact the regulation of RNA splicing (Esteve-Bruna et al. 2020; Payán-Bravo et al. 2021). These examples demonstrate the utility of uncovering RNA regulatory proteins via analysis of protein-protein interactions and suggest that other complexes present among Class 5 proteins without known roles in RNA regulation may warrant further investigation.

Complementary to the GO enrichment analysis, which inherently focused on Class 5 proteins that functioned together in particular pathways and complexes, we also examined individual Class 5 proteins that had few interactions with other Class 5 proteins but interacted with the Class 1, 2 and 3 protein complexes we identified earlier via clustering analysis (Figure 3C and 3D, Table S4). Because STRING transfers interaction data between homologous proteins from different species, this highlighted several proteins whose orthologs in other organisms are known RNA regulators but are not annotated as such in *C. elegans*. As examples, we focus on some of the Class 5 proteins with the highest number of interactions with Class 1, 2 and 3 proteins. For instance, *C. elegans* AATF-1 is only annotated as a transcriptional regulator, but our STRING network predicts interactions with 15 components of the small subunit processome (PPI cluster 3), an intermediate in the assembly of the small ribosomal subunit; indeed, the yeast and human AATF-1 orthologs have been found to function in ribosome biogenesis (Bernstein et al. 2004; Tafforeau et al. 2013; Badertscher et al. 2015; Kaiser et al. 2019; Singh et al. 2021). Similarly, NGP-1 is annotated only as a GTP binding protein, but is predicted by STRING to interact with several factors involved in biogenesis of the large ribosomal subunit (RPL-24.2 (part of PPI cluster 1 – cytoplasmic ribosome), NOG-1, and EIF-6); and again, the yeast and human orthologs of NGP-1 have been found to have roles in large subunit biogenesis (Matsuo et al. 2014; Liang et al. 2020). In fact, *C. elegans* NGP-1 has also been reported to play a role in nonsense-mediated mRNA decay (Casadio et al. 2015).

Other examples of Class 5 proteins identified in this manner with no currently annotated RNA regulatory roles in *C. elegans* include: PRMT-5, whose orthologs have roles in regulation of pre-mRNA splicing, cleavage and polyadenylation, and translation (Martin et al. 2010; Guderian et al. 2011; Bezzi et al. 2013; Lim et al. 2014; Gao et al. 2017), as reflected by the interactions observed in our network; and CYN-11 and MOA-2, both of which in our network are predicted to interact with several components of the spliceosome (PPI cluster 2), and are indeed orthologs of proteins (PPIH and SF3B5, respectively) that are known to be spliceosomal components in other organisms (Teigelkamp et al. 1998; Horowitz et al. 2002; Arribere et al. 2020).

Together, these analyses demonstrate the utility of our expanded RNA regulatory protein interaction network in uncovering additional putative co- and post-transcriptional regulators in *C. elegans* that are not currently identified in common annotation databases. While we have highlighted several examples here, our list of Class 5 proteins totalled 138, representing a substantial increase over the 1,353 proteins in Classes 1, 2 and 3. These will be interesting candidates to study in more detail, to elucidate any roles in RNA regulation in *C. elegans*. We do note, however, that the STRING database uses very low stringency criteria for defining orthologs, which could lead to erroneous transfer of interaction data between organisms. It will therefore be important to carefully assess the evidence for the predicted interactions of any given Class 5 protein when considering follow-up studies.

### Spatio-temporal expression of RNA regulatory genes suggests extensive opportunity to shape tissue development and function

To gain insight into how the RNA regulatory proteins that we have compiled might operate in the context of the development and function of *C. elegans* as a compact but complex multicellular organism, we analyzed data from several published studies describing tissue- and developmental stage-specific gene expression. Four datasets were examined: (1) single-cell RNA-seq data from L2 larvae (Cao et al. 2017), which provides transcript expression information across seven different tissues (gonad, intestine, body wall muscle, hypodermis, pharynx, glia, and neurons) as well as sub-tissue cell-type-specific information; (2) translating ribosome affinity purification (TRAP) coupled with RNA-seq in L4 larvae (Gracida et al. 2017), which provides transcript expression information at the level of ribosome association for intestine, body wall muscle, and neurons; (3) fluorescence-activated cell sorting (FACS) coupled to RNA-seq in adult animals (Kaletsky et al. 2018), which provides transcript expression data for intestine, body wall muscle, hypodermis, and neurons; and (4) FACS coupled to RNA-seq across 5 different timepoints of embryogenesis (Warner et al. 2019), with transcript expression information for intestine, muscle, hypodermis, pharynx, and neurons.

As a starting point we analyzed the single-cell RNA-seq data from Cao et al., 2017, using their analysis of transcript expression at the level of seven different tissues, since this was the largest number of tissues covered by any of the studies. We focused only on Class 1, 2, and 3 proteins, excluding Class 4, since the role of Class 4 proteins in RNA regulation is unclear, as discussed above. To look for common transcript expression patterns among different proteins, we performed k-means clustering, grouping 1,324 of the 1,353 Class 1-3 proteins into 13 different clusters (L2 expression clusters 1-13; Figure 4A and Table S2; see Materials and Methods for details). This revealed a variety of different expression patterns across tissues.

**Figure 4.**
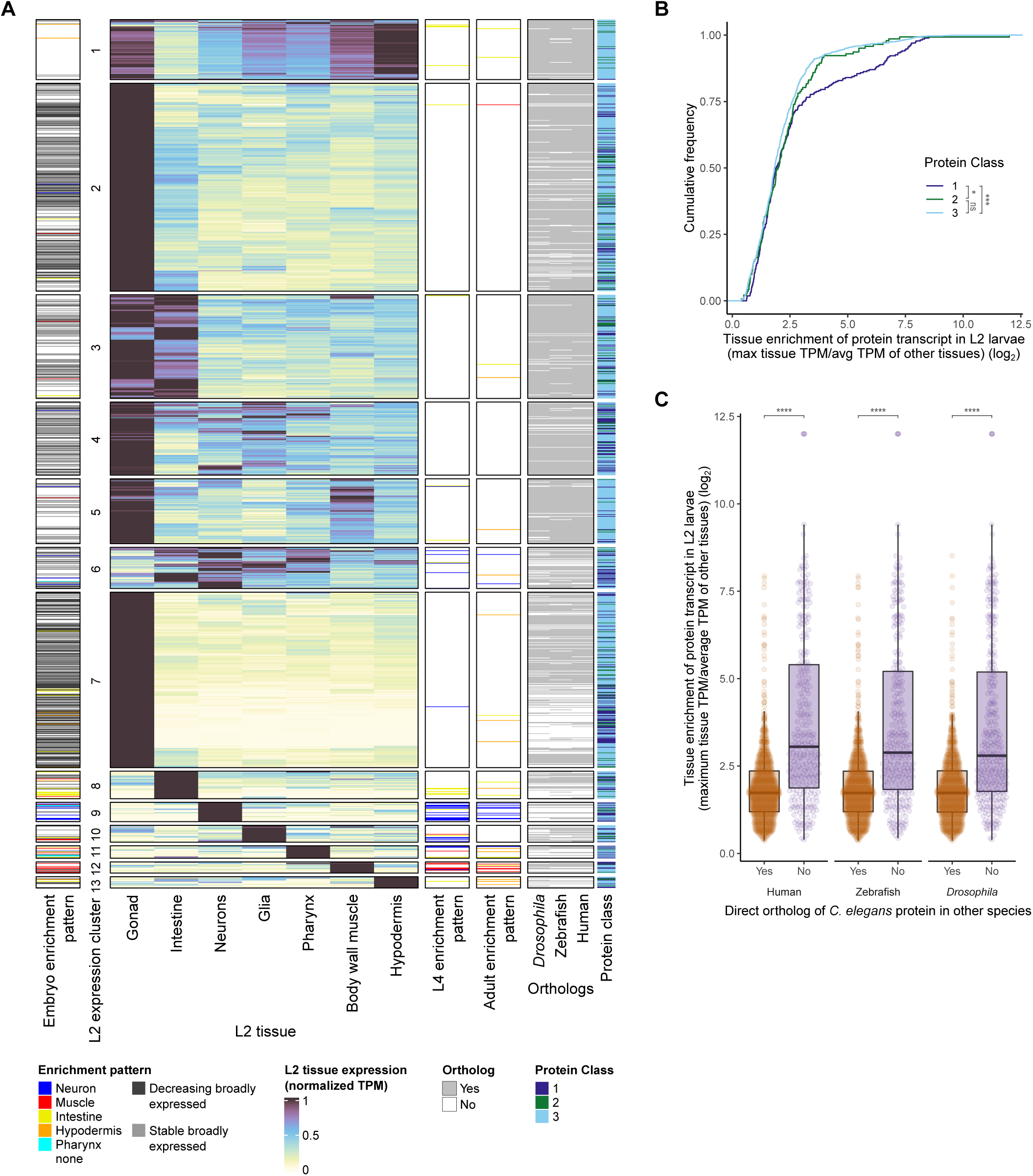
Overview of tissue- and developmental-stage-specific expression of Class 1, 2, and 3 RNA regulatory protein-encoding transcripts, as reported in the literature. **(A)** Expression of transcripts encoding RNA regulatory proteins from Classes 1, 2 and 3 in different tissues and stages of development. Rows correspond to individual RNA regulatory proteins and are ordered the same across all columns. Data shown are: (i) heatmap of tissue-specific normalized TPM calculated by Cao et al. 2017 for seven different tissues at the L2 larval stage; (ii) transcripts defined as enriched in various somatic tissues in the embryonic, L4 larval, and adult stages, by Warner et al. 2019, Gracida et al. 2017 and Kaletsky et al. 2018, respectively; embryo data also include annotation of transcripts that were broadly expressed across tissues and were either stable or decreasing in expression during the time-course of embryogenesis assessed by Warner et al.; (iii) the presence or absence of a direct ortholog of each *C. elegans* protein in *Drosophila melanogaster*, zebrafish, and humans; (iv) the corresponding RNA regulatory protein class. Proteins are grouped into 13 clusters by k-means clustering based on the L2 larval expression data. (B) Cumulative distributions, for RNA regulatory protein classes 1, 2, and 3, of the relative tissue specificity of their expression, in any given tissues, in the L2 larval data from Cao et al. 2017. Tissue specificity was calculated for each gene as the ratio of its highest TPM in any given tissue to the average TPM across the other six tissues. Statistical significance between distributions was calculated by two-sample Anderson-Darling test and is indicated: \**P* < 0.05; \*\*\**P* < 10^-5^; ns – not significant (*P* > 0.05). **(C)** Comparison of relative tissue specificity of expression, as measured in (B), between all Class 1, 2, and 3 *C. elegans* RNA regulatory proteins with (orange) or without (purple) a direct ortholog, in *Drosophila melanogaster*, zebrafish, or human. Statistical significance between proteins with and without a direct ortholog for each species was calculated by Wilcoxon rank sum test: \*\*\*\**P*< 10^-36^.

Most RNA regulatory protein genes in Classes 1, 2, and 3 were expressed in multiple tissues. Strikingly, however, the majority – 1,003 of 1,324 – were strongly expressed in the gonad, with varying degrees of expression elsewhere. Clusters with strong gonad expression included: cluster 2 (333 proteins), with strongest expression in the gonad, secondary expression in the intestine, and lower variable expression in most other tissues; cluster 3 (166 proteins), with strong, approximately equivalent expression in the gonad and intestine, and lower variable expression in most other tissues; cluster 4 (117 proteins), with strongest expression in the gonad and substantial secondary expression in most other non-intestinal tissues, particularly neurons, glia and pharynx; cluster 5 (105 proteins), with strongest expression in gonad, secondary expression in muscle, and lower variable expression in most other tissues; and cluster 7 (282 proteins), with predominant expression in gonad, and only much lower expression in any other tissues.

Outside of those clusters with strong gonadal expression, two showed broad expression across multiple tissues. Cluster 1 (97 proteins) had strong expression across tissues, and noticeably uniform expression for all proteins in the cluster; this cluster was composed almost entirely of cytoplasmic ribosomal proteins. Cluster 6 (66 proteins) had strong expression in neurons, intestine, glia and pharynx, weaker expression in hypodermis and muscle, and lowest expression in gonad.

The remaining proteins were divided into 6 clusters, each with predominant expression in a single somatic tissue: cluster 8 (44 proteins) in intestine; cluster 9 (31 proteins) in neurons; cluster 10 (27 proteins) in glia; cluster 11 (20 proteins) in pharynx; cluster 12 (18 proteins) in body wall muscle; and cluster 13 (18 proteins) in hypodermis.

Having established this detailed description of tissue expression patterns at the L2 larval stage, we next extended this analysis to later developmental stages by comparing the L2 larval expression data to tissue-specific expression data in L4 larvae obtained by TRAP-seq (Gracida et al. 2017), and in adults obtained by FACS coupled with RNA-seq (Kaletsky et al. 2018). As mentioned above, these latter datasets included expression data for intestine, neurons, muscle (L4 and adult) and hypodermis (adult only). Comparing genes identified in the respective studies as enriched for expression in one of the four somatic tissues in L4 or adults, to the cluster into which they were grouped based on our analysis of expression in L2 larvae, revealed a general agreement in the primary tissue of expression across these developmental stages, although this varied across tissues (Figure 4A and Table S2). For muscle, 8 of 11 proteins enriched in L4 muscle, and 6 of 7 proteins enriched in adult muscle, were part of L2 expression cluster 12 (predominantly muscle). For neurons, 13 of 27 proteins enriched in neurons at L4, and 8 of 11 proteins enriched in neurons in adults, were part of cluster 9 (predominantly neuronal), with another 5 of 27 in L4 and 2 of 11 in adult part of cluster 6 (strong expression in neurons, intestine, glia, and pharynx). For intestine, 5 of 19 proteins enriched in intestine at L4, and 2 of 8 proteins enriched in adult intestine, were part of cluster 8 (primarily intestine), with another 2 of 19 in L4 and 1 of 8 in adult being part of clusters 2 or 3 (strongest non-gonadal expression in intestine). For hypodermis, 4 of 13 proteins enriched in adult hypodermis were part of cluster 13 (primarily hypodermis).

Finally, we examined expression during embryogenesis, using tissue-specific expression data generated by FACS coupled to RNA-seq (Warner et al. 2019). For the five somatic tissues for which embryonic expression data are available – neurons, muscle, intestine, hypodermis, and pharynx – there is again general agreement with the expression patterns seen in L2 larvae (Figure 4A and Table S2). Based on embryonic tissue-enriched expression defined by Warner et al: 7 of 19 proteins enriched in embryonic muscle were part of L2 expression cluster 12 (predominantly muscle) and one was part of cluster 5 (gonad and muscle enriched); 7 of 17 proteins enriched in neurons in the embryo were part of cluster 9 (primarily neuron) and three were part of cluster 6 (strong expression in neurons, intestine, glia, and pharynx); 8 of 20 proteins enriched in intestine in the embryo were part of cluster 8 (predominantly intestine) and one was part of cluster 3 (strong expression in gonad and intestine); 2 of 5 proteins enriched in pharynx in the embryo were part of cluster 11 (predominantly pharynx); and 1 of 9 proteins enriched in hypodermis in the embryo was part of cluster 13 (primarily hypodermis).

Taken together, we can draw several conclusions from this analysis of RNA regulatory protein gene expression. First, the strong expression observed in the gonad in L2 larvae for the vast majority of these proteins suggests a particularly significant role for post-transcriptional regulation in this tissue, and is consistent with the established importance of post-transcriptional control in regulating the development of the germline (Nakamura and Seydoux 2008; Albarqi and Ryder 2023), which we explore in more detail below. Second, focusing on L2 larval expression in the six non-gonadal tissues, there is a significant degree of differential expression between tissues, and a substantial portion of RNA regulatory proteins – 158 in total – are expressed predominantly in just a single somatic tissue. These proteins may play important roles in sculpting the gene expression patterns required for tissue-specific development and functions. Finally, comparing patterns of tissue-enriched expression across developmental timepoints reveals a general agreement as to the predominant tissue of expression for any given protein at different stages, however we note that this was variable between tissues. While the significance of any apparent differences in tissue-enrichment across development for specific genes is difficult to interpret given the different laboratories and methods used to generate the datasets we examined, it is interesting that, in general, there was more agreement between muscle- and neuron-enriched expression across developmental stages, compared to intestine- and hypodermis-enriched genes. It’s possible that this might reflect a greater degree of developmental stage-specific expression of RNA regulatory proteins in the two latter tissues.

### Tissue-specificity of gene expression correlates with RNA regulatory protein class and evolutionary conservation

Given the varying levels of tissue-enrichment among RNA regulatory proteins, we sought to determine whether there were any features of the proteins that correlated with the degree of their tissue-specificity. First, we assessed whether there was any difference in the degree of tissue-specificity between proteins from the different RNA regulatory protein classes. To do this, we focussed on the L2 larval expression data and calculated, as a measure of tissue-specificity for each gene, the ratio of its expression in its most highly expressed tissue relative to the average of its expression in the remaining six tissues. Comparing the distribution of this measure between proteins in Classes 1, 2, and 3 revealed that a significantly higher proportion of Class 1 proteins were strongly tissue-enriched (Figure 4B). The difference in enrichment distributions was statistically significant comparing Class 1 versus 2 and Class 1 versus 3, but not Class 2 versus 3 (Two sample Anderson-Darling test *P*-values: Class 1 vs 2, *P* < 0.05; Class 1 vs 3, *P* < 10^-5^; Class 2 vs 3, *P* = 0.32). This suggests that Class 1 proteins have the most important roles in contributing to tissue-specific development and function. From a mechanistic view, this would be consistent with the fact that Class 1 proteins, through sequence-specific binding, have the potential to regulate expression of defined subsets of target RNAs, whereas most Class 2 and Class 3 proteins likely do not.

Second, we examined the evolutionary conservation of *C. elegans* RNA regulatory proteins in other model organism species, to determine whether there was any pattern of conservation correlating with the degree of tissue-specificity. For this analysis, we made use of data available at the Alliance of Genome Resources, which summarizes the predicted orthologs of genes across other model organisms, including *Drosophila*, zebrafish, and human. Using these data, we considered a *C. elegans* protein to have a direct ortholog in another species if it had at least one homolog in that species and was also the best match in *C. elegans* to the corresponding homolog; *C. elegans* proteins without a direct ortholog in another species therefore represent lineage-specific genes, derived through gene duplication (paralogs) or other mechanisms (Andersson et al. 2015). Comparing the tissue enrichment metric described above for proteins with or without a direct ortholog in human, zebrafish, or *Drosophila*, revealed that those proteins without direct orthologs showed significantly more tissue-specific expression patterns than those proteins with direct orthologs, for all three species comparisons (Figures 4A and 4C, and Table S2; Wilcoxon rank sum test *P* < 10^-36^). Although there are important exceptions to this trend, this suggests that, during its evolution, *C. elegans* has acquired unique post-transcriptional regulatory proteins which fulfill specialized tissue-specific functions. This is consistent with the general observation in *C. elegans* and other species that evolutionarily more recently duplicated genes show a higher degree of tissue-specific expression (Huminiecki and Wolfe 2004; Ma et al. 2024).

To highlight the RNA regulatory proteins most likely to contribute to tissue-specific development and function, we focussed on the Class 1 and 2 proteins that are classified as enriched in a specific somatic tissue in the embryonic, L2 larval, L4 larval, or adult stages. Figure S3 shows the expression of this set of proteins in all tissues at the L2 larval stage, as well as tissue enrichments described for the embryo, L4 larval, and adult stages, as above. We note that among the RNA regulatory proteins enriched in neurons across development are several post-transcriptional regulators with well-characterized roles in neurons, some of which are conserved across organisms, including *unc-75, ptb-1*, *mbl-1*, and *fox-1*, members of the mammalian CELF, PTBP, MBNL, and Rbfox families of RBPs, respectively.

### Gonad-enriched RNA regulatory proteins are expressed most strongly in the germline, and often exhibit tissue-specificity in their expression in non-gonadal somatic tissues

As noted earlier, the strong expression in the gonad of the vast majority of Class 1, 2 and 3 RNA regulatory proteins is likely a reflection of the importance of post-transcriptional control in regulating the development of the germline (Nakamura and Seydoux 2008; Albarqi and Ryder 2023). Interestingly, examining the embryonic expression patterns of these gonad-enriched proteins, described by Warner et al, 2019, we noticed that the proteins that were enriched in L2 larvae for gonad expression were largely classified by Warner et al.’s analysis as either “stable broadly expressed” or “decreasing broadly expressed” during the embryonic time-course that they assessed (Figure 4A and Table S2). This could be consistent with these being genes that are expressed in the germline, maternally during oogenesis, which then either have their expression maintained in embryos, or are degraded or diluted as development proceeds. To more directly evaluate whether these proteins are expressed in the germline, we examined the single-cell expression data from L2 larvae at a higher level of cell-type resolution, as reported by Cao et al., 2017. This analysis revealed that, among five gonadal cell types delineated in that study – germline, somatic gonad precursors, distal tip cells, vulval precursors, and sex myoblasts – 734 of 1,003 RNA regulatory genes falling into the clusters with strong gonad expression (clusters 2, 3, 4, 5, and 7) were most highly expressed in germline (Figure 5A), consistent with the idea that these gonad-enriched RNA regulatory proteins play an important role in germline development.

**Figure 5.**
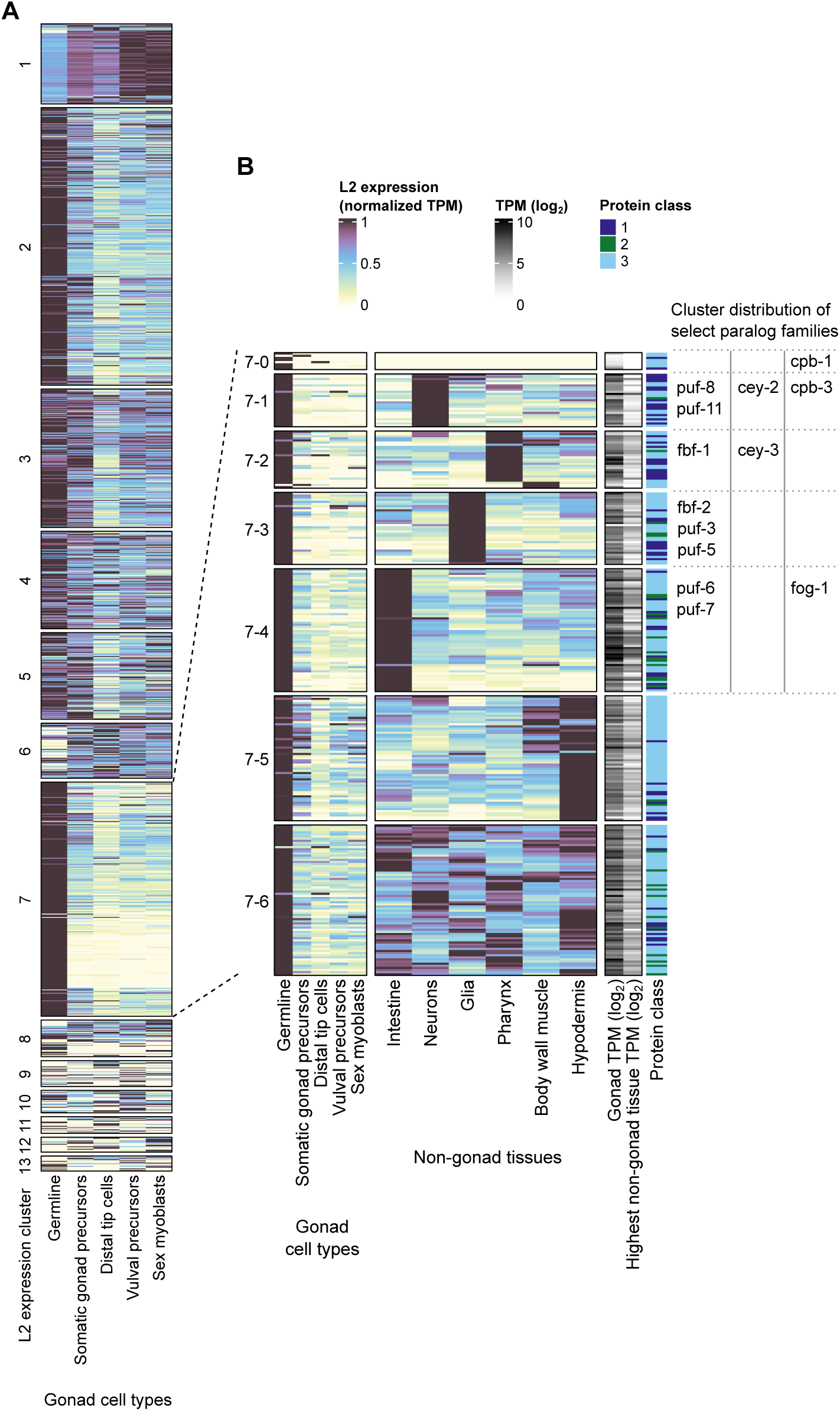
Analysis of the expression of gonad-enriched RNA regulatory proteins in gonad cell types and non-gonad somatic tissues in L2 larvae. **(A)** Analysis of Class 1-3 RNA regulatory protein expression in different gonad cell types, as reported by Cao et al., 2017. Clustering is as in Figure 4A, and data shown are TPM normalized to the highest gonad cell type TPM for each gene. **(B)** Expression of RNA regulatory genes in L2 expression cluster 7, in gonad cell types and non-gonad somatic tissues, as reported by Cao et al. 2017; gonad cell type expression data are normalized to the highest gonad cell type TPM for each gene, and non-gonad somatic tissue expression data are normalized to the highest non-gonad somatic tissue TPM for each gene. Since all L2 expression cluster 7 genes are highly enriched in the gonad compared to non-gonad somatic tissues, also shown are log_2_ TPM values for expression of each gene in gonad tissue and the highest non-gonad somatic tissue, for comparison. Genes are sub-clustered into 7 clusters based on normalized expression in non-gonad somatic tissues; note that the genes in cluster 7-0 did not have expression in non-gonad somatic tissues that passed our TPM cut-off (see Materials and Methods). To the right of the heatmaps are shown the distribution among the subclusters of members of three different protein paralog families – FBF/PUF, CEY, and CPB – highlighting their differential expression among non-gonad somatic tissues.

Interestingly, those genes that were most strongly enriched in the gonad relative to other tissues (L2 expression cluster 7) were also the most strongly enriched in the germline relative to other gonadal cell types (Figure 5A). Since the germline is an independent and particularly specialized cell lineage, we were curious to more closely examine the expression patterns of the germline-predominant RNA regulatory proteins in L2 expression cluster 7 in the other, non-gonad somatic tissues, to gain insight into any potential somatic functions of these proteins. While the expression of the L2 expression cluster 7 genes in non-gonadal tissues is generally substantially lower than their expression in gonad (median fold change of gonad TPM versus highest non-gonadal tissue TPM = 5.1; see also log_2_ TPM heatmap in Figure 5B), we note that the vast majority of these proteins are expressed in at least one non-gonadal L2 larval tissue. Of the 282 cluster 7 genes, eight did not have expression in any of the non-gonadal somatic tissues above our expression cut-off (see Materials and Methods), and we refer to these as subcluster 7-0. We performed k-means clustering on the remaining 274 cluster 7 proteins to group them into six clusters based on their normalized expression patterns in the non-gonadal somatic tissues; we refer to these clusters as L2 expression subclusters 7-1 through 7-6 (Figure 5B and Table S2). Strikingly, 144 of the 274 proteins showed predominant expression in a particular somatic tissue: subcluster 7-1 comprised 25 proteins predominantly expressed in neurons relative to the other somatic tissues; subcluster 7-2 comprised 27 proteins predominantly expressed in pharynx; subcluster 7-3 comprised 34 proteins predominantly expressed in glia; and subcluster 7-4 comprised 58 proteins predominantly expressed in intestine. A fifth subcluster, 7-5, showed strongest expression in hypodermis and muscle, but with substantial expression in the other somatic tissues as well. Subcluster 7-6 showed roughly equivalent expression across all somatic tissues. This suggests that in addition to their likely role in the germline, these proteins may have additional functions in specific somatic tissues.

Upon closer examination of the RNA regulatory proteins comprising these different subclusters, we noticed that for some protein families with multiple paralogs in *C. elegans*, the different paralogs were found in different subclusters, strongly enriched in different somatic tissues (Figure 5B). For example, eight of nine *C. elegans* PUF family proteins were part of L2 expression cluster 7, and these were distributed among subclusters 7-1, 7-2, 7-3, and 7-4, with predominant non-gonadal expression in neurons, pharynx, glia, or intestine, respectively (Figure 5B). Similarly, two of the four *C. elegans* Y-box-binding proteins were found in L2 expression cluster 7, and were differentially subclustered in 7-1 (neurons) and 7-2 (pharynx); and three of the four *C. elegans* CPEB family proteins were found in L2 expression cluster 7, and were divided into subclusters 7-0 (no substantial expression in non-gonadal tissue), 7-1 (neurons), and 7-4 (intestine). Interestingly, the genes from each of these families that were found in subcluster 7-1 with predominant non-gonadal expression in neurons – *puf-8*, *puf-11*, *cey-2*, and *cpb-3* – have all been independently shown either to be expressed in the nervous system or required for some aspect of nervous system development or function (Antonacci et al. 2015; R.N. Arey et al. 2019; R. Arey et al. 2019; Hayden et al. 2024), further supporting the potential functions of L2 expression cluster 7 proteins outside of the germline.

Taken together, these analyses support the idea that the predominance of RNA regulatory expression in the gonad is a reflection of the important role of post-transcriptional regulation in the germline. Moreover, our analysis shows that some RNA regulatory genes that are predominantly expressed in the germline are also expressed, albeit at lower levels, in specific non-gonadal somatic tissues. The differential somatic expression of paralogous protein family members that are all strongly expressed in the germline may offer insight into potential differences in their functions.

### Tissue-differentially expressed RNA regulatory proteins are enriched for different biological and molecular functions

To determine whether RNA regulatory proteins with different patterns of tissue-enriched expression are enriched for particular biological or molecular functions, we performed GO enrichment analysis for each of the 13 L2 larval expression clusters defined above, against the background of the set of all Class 1, 2, and 3 proteins for which L2 larval expression data were available in Cao et al. 2017. This revealed a variety of different GO terms enriched among the different expression clusters (Figure 6A and Table S6).

**Figure 6.**
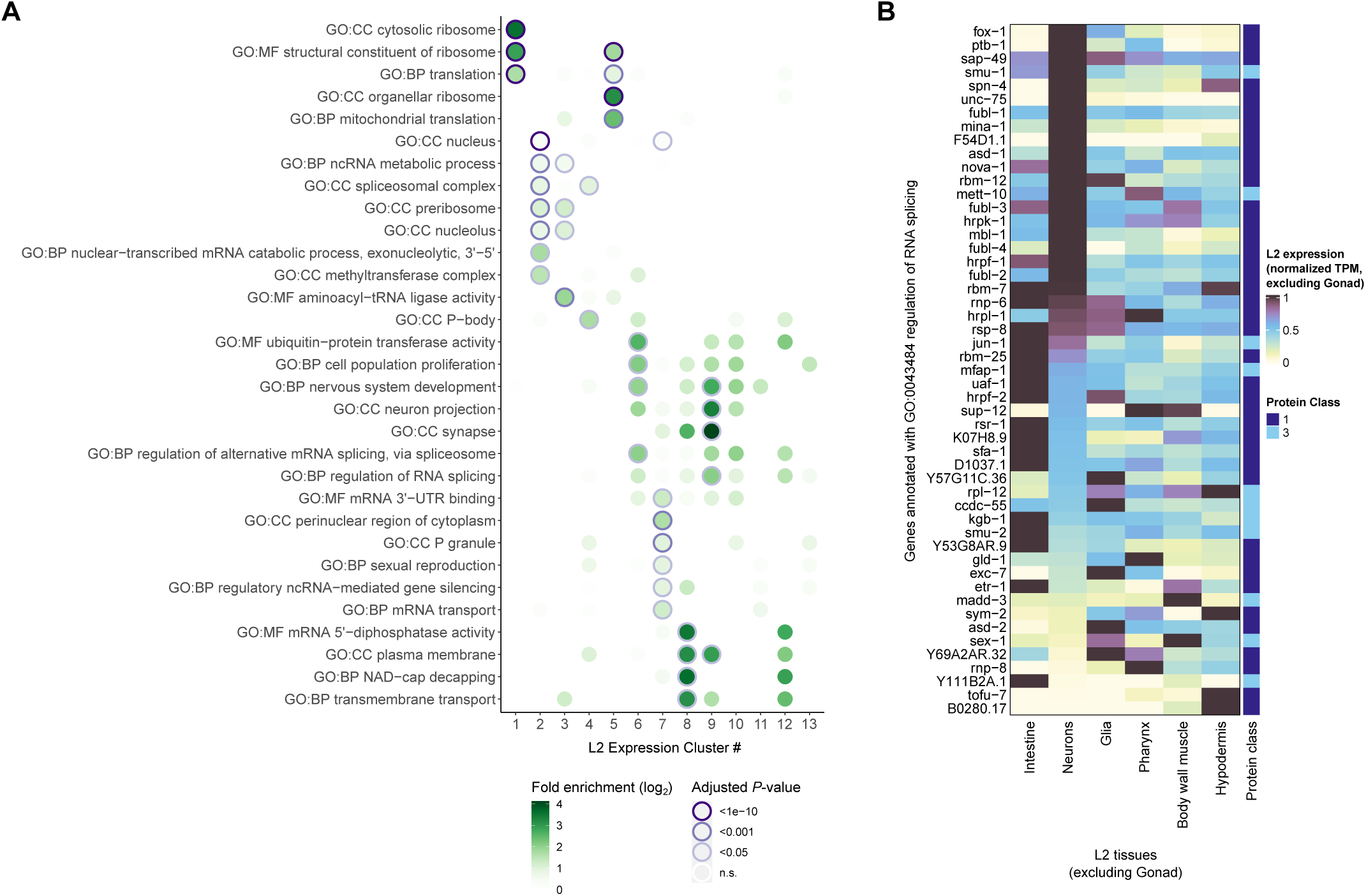
Functional enrichments among Class 1-3 RNA regulatory protein clusters with different patterns of tissue-specific expression in L2 larvae. **(A)** Summary of GO term enrichment analysis performed on each of the L2 larval expression clusters, against the background of all Class 1, 2, and 3 RNA regulatory proteins. Fold enrichment and adjusted *P*-value for each GO term/cluster pairing are indicated by the fill and outline of the circles, respectively. Cluster numbering is as in Figure 4A. Full results of the GO enrichment analysis are presented in Table S6. **(B)** Heatmap showing tissue-specific expression of all *C. elegans* genes annotated with the GO term “regulation of RNA splicing” (GO:0043484), in L2 larvae for the six non-gonad tissues reported by Cao et al., 2017. Expression values shown for each gene are normalized to the highest TPM among the six non-gonad tissues. The RNA regulatory protein class to which each gene belongs is indicated on the right.

For some clusters, a subset of the enriched GO terms reflected the function or characteristics of the tissues in which their genes were predominantly expressed. For example, cluster 6 (strong expression in neurons, glia, pharynx, and intestine) and cluster 9 (predominantly expressed in neurons) were enriched for terms corresponding to nervous system development and function, whereas cluster 7 (predominantly gonad expression) was enriched for the GO term “sexual reproduction.”

GO term analysis also revealed that several post-transcriptional functions or complexes were enriched in specific L2 expression clusters. For example, cluster 2 genes (strong gonad expression and lower variable expression across other tissues) were enriched for GO terms describing the spliceosome and the pre-ribosome; cluster 7 genes (predominant gonad expression) were enriched for components of P granules, a class of germline-specific RNA granule in *C. elegans*; cluster 1 genes (strong expression across all tissues) were enriched for GO terms describing the cytoplasmic ribosome; and cluster 5 genes (strong gonad and broad secondary expression in somatic tissues, highest in muscle) were enriched for GO terms describing the mitochondrial ribosome. The enrichment for cytoplasmic ribosome components among cluster 1 proteins is consistent with the ubiquitous requirement of the ribosome in all tissues. Likewise, the enrichment for mitochondrial ribosome components among cluster 5 proteins may reflect a broad requirement for mitochondrial translation across tissues, but with higher mitochondrial demands in muscle.

We were particularly interested to note that GO terms describing the regulation of splicing and alternative splicing were enriched in the two clusters characterized by strong neuronal expression, and which were also enriched for GO terms related to nervous system development: cluster 6 (strong expression in neurons, glia, pharynx, and intestine) and cluster 9 (predominantly neuronal expression). To gain a more complete picture of the expression of splicing regulatory factors across somatic tissues in L2 larvae, we calculated expression values normalized across the six non-gonad tissues reported by Cao et al. 2017 (intestine, neurons, glia, pharynx, muscle, and hypodermis), for all *C. elegans* genes annotated with the GO term “regulation of RNA splicing” (Figure 6B). Strikingly, nearly half of these putative splicing regulators had higher expression in neurons than in other somatic tissues. This is consistent with the observation in *C. elegans* and other organisms that neurons exhibit higher levels of alternative splicing than other somatic tissues (Barbosa-Morais et al. 2012; Tapial et al. 2017; Koterniak et al. 2020), which recent work in *C. elegans* has revealed is likely due to differential alternative splicing among neuronal subtypes (Weinreb et al. 2024; Wolfe et al. 2024); this increased complexity of alternative splicing would be expected to require a more diverse set of splicing regulators. We expect that our analysis of splicing regulator expression in L2 larval tissues will help to guide further studies aimed at understanding the control of alternative splicing in neurons.

## Conclusion

Here, we have provided a global overview of the RNA regulatory protein landscape in *C. elegans*. We compiled a comprehensive catalogue of 2,152 RNA regulatory proteins, including canonical sequence- or structure-specific RNA-binding proteins (283 proteins, Class 1), non-specific RNA-binding proteins (144 proteins, Class 2), indirect RNA-associated proteins or post-translational regulators annotated as having roles in RNA regulation or function (926 proteins, Class 3), and proteins identified as binding to RNA based on RNA interactome capture experiments in *C. elegans* (Matia-González et al. 2015; Esmaillie et al. 2019) (799 proteins, Class 4). We used available protein interaction data in the STRING database to create a putative protein-protein interaction network highlighting the connections between these RNA regulatory proteins, and leveraged this network to identify 138 additional proteins with likely roles in RNA regulatory processes that were previously unannotated. Lastly, we analyzed tissue- and developmental stage-specific expression data from several published studies, revealing tissue-enriched expression for a majority of RNA regulatory proteins. This highlighted the particularly important role of post-transcriptional regulatory processes in the germline, and their potential for sculpting the development and function of individual somatic tissues. Together, this atlas of RNA regulatory proteins, their interactions, and their expression patterns, will provide a useful resource for future studies of *C. elegans* RNA biology, and a template for the compilation of similar atlases in other species. More broadly, it provides detailed insight into the global organization of an RNA regulatory protein network in a multicellular organism.

## Data Availability

The authors affirm that all data and analyses generated in this study are represented fully within the article, its figures, and its supplemental tables, figures and files. Files S1 and S3 are available at: https://figshare.com/articles/dataset/Laver_et_al_2024_FilesS1_and_S3/27629376. Previously published datasets used in this study were from: Matia-González et al. 2015, Cao et al. 2017, Gracida et al. 2017, Kaletsky et al. 2018, Esmaillie et al. 2019, Warner et al. 2019, the STRING database (https://string-db.org/), and the Alliance of Genome Resources (https://www.alliancegenome.org/), as described in the Materials and Methods.

## Author Contributions

JAC and JDL conceptualized the study. JDL performed the analyses, prepared the figures, and wrote the manuscript. MD wrote the initial draft of the manuscript introduction and performed preliminary protein-protein interaction analyses. JAC provided supervision to JDL and MD, and provided critical feedback on the analyses, figures, and manuscript.

## Acknowledgments

We thank members of the Calarco Lab for valuable suggestions and feedback on analyses and presentation of data. JDL was funded in part by a CIHR Fellowship, with additional funds from the University of Toronto. This research was supported by CIHR project grants and an NSERC Discovery Grant to J.A.C.

## Conflicts of Interest

The authors declare no conflicts of interest.

## List of Supplemental Materials

**Table S1.** Lists of sequence-specific RNA-binding domains, non-specific RNA-binding and metabolism-related domains, Gene Ontology terms, and UniProt keywords used to compile complete list of *C. elegans* RNA regulatory proteins.

**Table S2.** The *C. elegans* RNA regulatory proteome: complete list of all RNA regulatory proteins in Classes 1, 2, 3, and 4. For each protein, includes: (i) WBGeneID; (ii) sequence ID; (iii) gene name; (iv) protein class as defined in this study; (v) whether contains a sequence- or structure-specific RBD, and associated IPR identifier; (vi) whether contains a non-specific RBD or RNA metabolism-related domain, and associated IPR identifier; (vii) annotated with RNA regulation-related GO term (Yes/No); (viii) annotated with RNA regulation-related UniProt keyword (Yes/No); (ix) included in RBP list from Tamburino et al. 2013 (Yes/No); (x) identified in RNA interactome capture studies (Yes/No); (xi) protein-protein interaction network clustering result; (xii) summary of expression data used in analyses; (xiii) presence of direct orthologs in *Drosophila*, zebrafish, and human.

**Table S3.** Full GO term enrichment results for PPI clusters 1-22. Relevant to Figure 2.

**Table S4.** List of Class 5 proteins, with number of interactions to Class 1-3 proteins, and to other Class 5 proteins. Relevant to Figure 3.

**Table S5.** Full GO term enrichment results for Class 5 proteins. Relevant to Figure 3.

**Table S6.** Full GO term enrichment results for L2 larvae expression clusters 1-13. Relevant to Figure 6.

**Figure S1.** Full high-resolution image of complete RNA regulatory protein-protein interaction (Classes 1-4), with nodes labelled with gene names. Relevant to Figure 2.

**Figure S2.** Full high-resolution image of expanded RNA regulatory protein-protein interaction (Classes 1, 2, 3, 5), with nodes labelled with gene names. Relevant to Figure 3.

**Figure S3.** Overview of highly tissue-enriched RNA regulatory proteins in Classes 1 and 2.

**File S1.** Cytoscape-loadable protein-protein interaction network file (.cys) for complete RNA regulatory protein interaction network (Classes 1-4). Relevant to Figure 2. Available at: https://figshare.com/articles/dataset/Laver_et_al_2024_FilesS1_and_S3/27629376.

**File S2.** Protein-protein interaction network file (.sif) for full RNA regulatory protein list, Classes 1-4. Relevant to Figure 2.

**File S3.** Cytoscape-loadable protein-protein interaction network file (.cys) for expanded RNA regulatory protein interaction network (Classes 1, 2, 3, 5). Relevant to Figure 3. Available at: https://figshare.com/articles/dataset/Laver_et_al_2024_FilesS1_and_S3/27629376.

**File S4.** Protein-protein interaction network file (.sif) for proteins in Classes 1, 2, 3 and 5. Relevant to Figure 3.

## Notes

https://figshare.com/articles/dataset/Laver_et_al_2024_FilesS1_and_S3/27629376

### Competing Interest Statement

The authors have declared no competing interest.

## References

Aguilera-Gomez A, Zacharogianni M, Oorschot MM van, Genau H, Grond R, Veenendaal T, Sinsimer KS, Gavis ER, Behrends C, Rabouille C. 2017. Phospho-Rasputin Stabilization by Sec16 Is Required for Stress Granule Formation upon Amino Acid Starvation. Cell Reports. 20(4):935–948. doi:10.1016/j.celrep.2017.06.042.

Albarqi MMY, Ryder SP. 2023. The role of RNA-binding proteins in orchestrating germline development in Caenorhabditis elegans. Frontiers in Cell and Developmental Biology. 10. [accessed 2023 Jan 23]. https://www.frontiersin.org/articles/10.3389/fcell.2022.1094295.

Alliance of Genome Resources Consortium. 2024. Updates to the Alliance of Genome Resources central infrastructure. Genetics. 227(1):iyae049. doi:10.1093/genetics/iyae049.

Andersson DI, Jerlström-Hultqvist J, Näsvall J. 2015. Evolution of New Functions De Novo and from Preexisting Genes. Cold Spring Harb Perspect Biol. 7(6):a017996. doi:10.1101/cshperspect.a017996.

Antonacci S, Forand D, Wolf M, Tyus C, Barney J, Kellogg L, Simon MA, Kerr G, Wells KL, Younes S, et al. 2015. Conserved RNA-Binding Proteins Required for Dendrite Morphogenesis in Caenorhabditis elegans Sensory Neurons. G3 Genes|Genomes|Genetics. 5(4):639–653. doi:10.1534/g3.115.017327.

Arey R, Kaletsky R, Murphy CT. 2019. Profiling of presynaptic mRNAs reveals a role for axonal PUMILIOs in associative memory formation. :733428. doi:10.1101/733428. [accessed 2024 Sep 20]. https://www.biorxiv.org/content/10.1101/733428v1.

Arey RN, Kaletsky R, Murphy CT. 2019. Nervous system-wide profiling of presynaptic mRNAs reveals regulators of associative memory. Sci Rep. 9(1):20314. doi:10.1038/s41598-019-56908-8.

Arribere JA, Kuroyanagi H, Hundley HA. 2020. mRNA Editing, Processing and Quality Control in Caenorhabditis elegans. Genetics. 215(3):531–568. doi:10.1534/genetics.119.301807.

Bader AS, Hawley BR, Wilczynska A, Bushell M. 2020. The roles of RNA in DNA double-strand break repair. Br J Cancer. 122(5):613–623. doi:10.1038/s41416-019-0624-1.

Badertscher L, Wild T, Montellese C, Alexander LT, Bammert L, Sarazova M, Stebler M, Csucs G, Mayer TU, Zamboni N, et al. 2015. Genome-wide RNAi Screening Identifies Protein Modules Required for 40S Subunit Synthesis in Human Cells. Cell Reports. 13(12):2879–2891. doi:10.1016/j.celrep.2015.11.061.

Baltz AG, Munschauer M, Schwanhäusser B, Vasile A, Murakawa Y, Schueler M, Youngs N, Penfold-Brown D, Drew K, Milek M, et al. 2012. The mRNA-bound proteome and its global occupancy profile on protein-coding transcripts. Mol Cell. 46(5):674–690. doi:10.1016/j.molcel.2012.05.021.

Barbet NC, Schneider U, Helliwell SB, Stansfield I, Tuite MF, Hall MN. 1996. TOR controls translation initiation and early G1 progression in yeast. Molecular Biology of the Cell. 7(1):25–42. doi:10.1091/mbc.7.1.25.

Barbosa-Morais NL, Irimia M, Pan Q, Xiong HY, Gueroussov S, Lee LJ, Slobodeniuc V, Kutter C, Watt S, Çolak R, et al. 2012. The Evolutionary Landscape of Alternative Splicing in Vertebrate Species. Science. 338(6114):1587–1593. doi:10.1126/science.1230612.

Baßler J, Hurt E. 2019. Eukaryotic Ribosome Assembly. Annu Rev Biochem. 88:281–306. doi:10.1146/annurev-biochem-013118-110817.

Berkyurek AC, Furlan G, Lampersberger L, Beltran T, Weick E-M, Nischwitz E, Cunha Navarro I, Braukmann F, Akay A, Price J, et al. 2021. The RNA polymerase II subunit RPB-9 recruits the integrator complex to terminate Caenorhabditis elegans piRNA transcription. EMBO J. 40(5):e105565. doi:10.15252/embj.2020105565.

Bernstein KA, Gallagher JEG, Mitchell BM, Granneman S, Baserga SJ. 2004. The Small-Subunit Processome Is a Ribosome Assembly Intermediate. Eukaryotic Cell. 3(6):1619–1626. doi:10.1128/ec.3.6.1619-1626.2004.

Bezzi M, Teo SX, Muller J, Mok WC, Sahu SK, Vardy LA, Bonday ZQ, Guccione E. 2013. Regulation of constitutive and alternative splicing by PRMT5 reveals a role for Mdm4 pre-mRNA in sensing defects in the spliceosomal machinery. Genes Dev. 27(17):1903–1916. doi:10.1101/gad.219899.113.

Boreikaitė V, Passmore LA. 2023. 3’-End Processing of Eukaryotic mRNA: Machinery, Regulation, and Impact on Gene Expression. Annu Rev Biochem. 92:199–225. doi:10.1146/annurev-biochem-052521-012445.

Bourke AM, Schwarz A, Schuman EM. 2023. De-centralizing the Central Dogma: mRNA translation in space and time. Molecular Cell. 83(3):452–468. doi:10.1016/j.molcel.2022.12.030.

Bovaird S, Patel D, Padilla J-CA, Lécuyer E. 2018. Biological functions, regulatory mechanisms, and disease relevance of RNA localization pathways. FEBS Letters. 592(17):2948–2972. doi:10.1002/1873-3468.13228.

Brunn GJ, Hudson CC, Sekulić A, Williams JM, Hosoi H, Houghton PJ, Jr. JCL, Abraham RT. 1997. Phosphorylation of the Translational Repressor PHAS-I by the Mammalian Target of Rapamycin. Science. 277(5322):99–101. doi:10.1126/science.277.5322.99.

Cao J, Packer JS, Ramani V, Cusanovich DA, Huynh C, Daza R, Qiu X, Lee C, Furlan SN, Steemers FJ, et al. 2017. Comprehensive single-cell transcriptional profiling of a multicellular organism. Science. 357(6352):661–667. doi:10.1126/science.aam8940.

Casadio A, Longman D, Hug N, Delavaine L, Vallejos Baier R, Alonso CR, Cáceres JF. 2015. Identification and characterization of novel factors that act in the nonsense-mediated mRNA decay pathway in nematodes, flies and mammals. EMBO reports. 16(1):71–78. doi:10.15252/embr.201439183.

Castello A, Fischer B, Eichelbaum K, Horos R, Beckmann BM, Strein C, Davey NE, Humphreys DT, Preiss T, Steinmetz LM, et al. 2012. Insights into RNA biology from an atlas of mammalian mRNA-binding proteins. Cell. 149(6):1393–1406. doi:10.1016/j.cell.2012.04.031.

Castello A, Fischer B, Frese CK, Horos R, Alleaume A-M, Foehr S, Curk T, Krijgsveld J, Hentze MW. 2016. Comprehensive Identification of RNA-Binding Domains in Human Cells. Molecular Cell. 63(4):696–710. doi:10.1016/j.molcel.2016.06.029.

Choder M. 2004. Rpb4 and Rpb7: subunits of RNA polymerase II and beyond. Trends in Biochemical Sciences. 29(12):674–681. doi:10.1016/j.tibs.2004.10.007.

Cook KB, Kazan H, Zuberi K, Morris Q, Hughes TR. 2011. RBPDB: a database of RNA-binding specificities. Nucleic Acids Research. 39(suppl_1):D301–D308. doi:10.1093/nar/gkq1069.

Cookson MR. 2017. RNA-binding proteins implicated in neurodegenerative diseases. WIREs RNA. 8(1):e1397. doi:10.1002/wrna.1397.

Corley M, Burns MC, Yeo GW. 2020. How RNA-Binding Proteins Interact with RNA: Molecules and Mechanisms. Molecular Cell. 78(1):9–29. doi:10.1016/j.molcel.2020.03.011.

Dahan N, Choder M. 2013. The eukaryotic transcriptional machinery regulates mRNA translation and decay in the cytoplasm. Biochimica et Biophysica Acta (BBA) - Gene Regulatory Mechanisms. 1829(1):169–173. doi:10.1016/j.bbagrm.2012.08.004.

Das S, Vera M, Gandin V, Singer RH, Tutucci E. 2021. Intracellular mRNA transport and localized translation. Nat Rev Mol Cell Biol. 22(7):483–504. doi:10.1038/s41580-021-00356-8.

Doncheva NT, Morris JH, Gorodkin J, Jensen LJ. 2019. Cytoscape StringApp: Network Analysis and Visualization of Proteomics Data. J Proteome Res. 18(2):623–632. doi:10.1021/acs.jproteome.8b00702.

Doncheva NT, Morris JH, Holze H, Kirsch R, Nastou KC, Cuesta-Astroz Y, Rattei T, Szklarczyk D, von Mering C, Jensen LJ. 2023. Cytoscape stringApp 2.0: Analysis and Visualization of Heterogeneous Biological Networks. J Proteome Res. 22(2):637–646. doi:10.1021/acs.jproteome.2c00651.

Dowd C. 2023. twosamples: Fast Permutation Based Two Sample Tests. https://twosampletest.com.

Esmaillie R, Ignarski M, Bohl K, Krüger T, Ahmad D, Seufert L, Schermer B, Benzing T, Müller R-U, Fabretti F. 2019. Activation of Hypoxia-Inducible Factor Signaling Modulates the RNA Protein Interactome in Caenorhabditis elegans. iScience. 22:466–476. doi:10.1016/j.isci.2019.11.039.

Esteve-Bruna D, Carrasco-López C, Blanco-Touriñán N, Iserte J, Calleja-Cabrera J, Perea-Resa C, Úrbez C, Carrasco P, Yanovsky MJ, Blázquez MA, et al. 2020. Prefoldins contribute to maintaining the levels of the spliceosome LSM2–8 complex through Hsp90 in Arabidopsis. Nucleic Acids Research. 48(11):6280–6293. doi:10.1093/nar/gkaa354.

Gao G, Dhar S, Bedford MT. 2017. PRMT5 regulates IRES-dependent translation via methylation of hnRNP A1. Nucleic Acids Research. 45(8):4359–4369. doi:10.1093/nar/gkw1367.

Gerstberger S, Hafner M, Tuschl T. 2014. A census of human RNA-binding proteins. Nat Rev Genet. 15(12):829–845. doi:10.1038/nrg3813.

Gracida X, Dion MF, Harris G, Zhang Y, Calarco JA. 2017. An Elongin-Cullin-SOCS Box Complex Regulates Stress-Induced Serotonergic Neuromodulation. Cell Reports. 21(11):3089– 3101. doi:10.1016/j.celrep.2017.11.042.

Gu Z. 2022. Complex heatmap visualization. iMeta. 1(3):e43. doi:10.1002/imt2.43.

Gu Z, Eils R, Schlesner M. 2016. Complex heatmaps reveal patterns and correlations in multidimensional genomic data. Bioinformatics. 32(18):2847–2849. doi:10.1093/bioinformatics/btw313.

Guderian G, Peter C, Wiesner J, Sickmann A, Schulze-Osthoff K, Fischer U, Grimmler M. 2011. RioK1, a New Interactor of Protein Arginine Methyltransferase 5 (PRMT5), Competes with pICln for Binding and Modulates PRMT5 Complex Composition and Substrate Specificity. J Biol Chem. 286(3):1976–1986. doi:10.1074/jbc.M110.148486.

Hagerman PJ, Hagerman RJ. 2015. Fragile X–associated tremor/ataxia syndrome. Annals of the New York Academy of Sciences. 1338(1):58–70. doi:10.1111/nyas.12693.

Hardiman O, Al-Chalabi A, Chio A, Corr EM, Logroscino G, Robberecht W, Shaw PJ, Simmons Z, van den Berg LH. 2017. Amyotrophic lateral sclerosis. Nat Rev Dis Primers. 3(1):1–19. doi:10.1038/nrdp.2017.71.

Hayden AN, Brandel KL, Merlau PR, Vijayakumar P, Leptich EJ, Pietryk EW, Gaytan ES, Ni CW, Chao H-T, Rosenfeld JA, et al. 2024. Behavioral screening of conserved RNA-binding proteins reveals CEY-1/YBX RNA-binding protein dysfunction leads to impairments in memory and cognition. :2024.01.05.574402. doi:10.1101/2024.01.05.574402. [accessed 2024 Sep 20]. https://www.biorxiv.org/content/10.1101/2024.01.05.574402v2.

Heck AM, Wilusz J. 2018. The Interplay between the RNA Decay and Translation Machinery in Eukaryotes. Cold Spring Harb Perspect Biol. 10(5):a032839. doi:10.1101/cshperspect.a032839.

Hentze MW, Castello A, Schwarzl T, Preiss T. 2018. A brave new world of RNA-binding proteins. Nat Rev Mol Cell Biol. 19(5):327–341. doi:10.1038/nrm.2017.130.

Hofweber M, Dormann D. 2019. Friend or foe—Post-translational modifications as regulators of phase separation and RNP granule dynamics. Journal of Biological Chemistry. 294(18):7137– 7150. doi:10.1074/jbc.TM118.001189.

Horos R, Büscher M, Kleinendorst R, Alleaume A-M, Tarafder AK, Schwarzl T, Dziuba D, Tischer C, Zielonka EM, Adak A, et al. 2019. The Small Non-coding Vault RNA1-1 Acts as a Riboregulator of Autophagy. Cell. 176(5):1054–1067.e12. doi:10.1016/j.cell.2019.01.030.

Horowitz DS, Lee EJ, Mabon SA, Misteli T. 2002. A cyclophilin functions in pre-mRNA splicing. The EMBO Journal. 21(3):470–480. doi:10.1093/emboj/21.3.470.

Hu G, McQuiston T, Bernard A, Park Y-D, Qiu J, Vural A, Zhang N, Waterman SR, Blewett NH, Myers TG, et al. 2015. A conserved mechanism of TOR-dependent RCK-mediated mRNA degradation regulates autophagy. Nature Cell Biology. 17(7):930–942. doi:10.1038/ncb3189.

Huminiecki L, Wolfe KH. 2004. Divergence of Spatial Gene Expression Profiles Following Species-Specific Gene Duplications in Human and Mouse. Genome Res. 14(10a):1870–1879. doi:10.1101/gr.2705204.

Huppertz I, Perez-Perri JI, Mantas P, Sekaran T, Schwarzl T, Russo F, Ferring-Appel D, Koskova Z, Dimitrova-Paternoga L, Kafkia E, et al. 2022. Riboregulation of Enolase 1 activity controls glycolysis and embryonic stem cell differentiation. Molecular Cell. 82(14):2666–2680.e11. doi:10.1016/j.molcel.2022.05.019.

Jefferies HBJ, Fumagalli S, Dennis PB, Reinhard C, Pearson RB, Thomas G. 1997. Rapamycin suppresses 5′TOP mRNA translation through inhibition of p70s6k. The EMBO Journal. 16(12):3693–3704. doi:10.1093/emboj/16.12.3693.

Kaiser RWJ, Ignarski M, Van Nostrand EL, Frese CK, Jain M, Cukoski S, Heinen H, Schaechter M, Seufert L, Bunte K, et al. 2019. A protein-RNA interaction atlas of the ribosome biogenesis factor AATF. Sci Rep. 9(1):11071. doi:10.1038/s41598-019-47552-3.

Kaletsky R, Yao V, Williams A, Runnels AM, Tadych A, Zhou S, Troyanskaya OG, Murphy CT. 2018. Transcriptome analysis of adult Caenorhabditis elegans cells reveals tissue-specific gene and isoform expression. PLOS Genetics. 14(8):e1007559. doi:10.1371/journal.pgen.1007559.

Karousis ED, Mühlemann O. 2019. Nonsense-Mediated mRNA Decay Begins Where Translation Ends. Cold Spring Harb Perspect Biol. 11(2):a032862. doi:10.1101/cshperspect.a032862.

Khanduja JS, Calvo IA, Joh RI, Hill IT, Motamedi M. 2016. Nuclear Noncoding RNAs and Genome Stability. Molecular Cell. 63(1):7–20. doi:10.1016/j.molcel.2016.06.011.

Kolberg L, Raudvere U, Kuzmin I, Adler P, Vilo J, Peterson H. 2023. g:Profiler—interoperable web service for functional enrichment analysis and gene identifier mapping (2023 update). Nucleic Acids Research. 51(W1):W207–W212. doi:10.1093/nar/gkad347.

Koterniak B, Pilaka PP, Gracida X, Schneider L-M, Pritišanac I, Zhang Y, Calarco JA. 2020. Global regulatory features of alternative splicing across tissues and within the nervous system of C. elegans. Genome Res. 30(12):1766–1780. doi:10.1101/gr.267328.120.

Lee JE, Cathey PI, Wu H, Parker R, Voeltz GK. 2020. Endoplasmic reticulum contact sites regulate the dynamics of membraneless organelles. Science. 367(6477):eaay7108. doi:10.1126/science.aay7108.

Liang X, Zuo M-Q, Zhang Y, Li N, Ma C, Dong M-Q, Gao N. 2020. Structural snapshots of human pre-60S ribosomal particles before and after nuclear export. Nat Commun. 11(1):3542. doi:10.1038/s41467-020-17237-x.

Lim J-H, Lee Y-M, Lee G, Choi Y-J, Lim B-O, Kim YJ, Choi D-K, Park J-W. 2014. PRMT5 is essential for the eIF4E-mediated 5′-cap dependent translation. Biochemical and Biophysical Research Communications. 452(4):1016–1021. doi:10.1016/j.bbrc.2014.09.033.

Loedige I, Stotz M, Qamar S, Kramer K, Hennig J, Schubert T, Löffler P, Längst G, Merkl R, Urlaub H, et al. 2014. The NHL domain of BRAT is an RNA-binding domain that directly contacts the hunchback mRNA for regulation. Genes Dev. 28(7):749–764. doi:10.1101/gad.236513.113.

Ma F, Lau CY, Zheng C. 2024. Young duplicate genes show developmental stage- and cell type-specific expression and function in *Caenorhabditis elegans*. Cell Genomics. 4(1):100467. doi:10.1016/j.xgen.2023.100467.

Machado de Amorim A, Chakrabarti S. 2022. Assembly of multicomponent machines in RNA metabolism: A common theme in mRNA decay pathways. WIREs RNA. 13(2):e1684. doi:10.1002/wrna.1684.

Martin G, Ostareck-Lederer A, Chari A, Neuenkirchen N, Dettwiler S, Blank D, Rüegsegger U, Fischer U, Keller W. 2010. Arginine methylation in subunits of mammalian pre-mRNA cleavage factor I. RNA. 16(8):1646–1659. doi:10.1261/rna.2164210.

Matia-González AM, Laing EE, Gerber AP. 2015. Conserved mRNA-binding proteomes in eukaryotic organisms. Nature Structural & Molecular Biology. 22(12):1027–1033. doi:10.1038/nsmb.3128.

Matsuo Y, Granneman S, Thoms M, Manikas R-G, Tollervey D, Hurt E. 2014. Coupled GTPase and remodelling ATPase activities form a checkpoint for ribosome export. Nature. 505(7481):112–116. doi:10.1038/nature12731.

Mercuri E, Sumner CJ, Muntoni F, Darras BT, Finkel RS. 2022. Spinal muscular atrophy. Nat Rev Dis Primers. 8(1):1–16. doi:10.1038/s41572-022-00380-8.

Milano SN, Bayer LV, Ko JJ, Casella CE, Bratu DP. 2024. The role of ER exit sites in maintaining P-body organization and transmitting ER stress response during Drosophila melanogaster oogenesis. :2024.07.03.601952. doi:10.1101/2024.07.03.601952. [accessed 2024 Jul 8]. https://www.biorxiv.org/content/10.1101/2024.07.03.601952v1.

Moore MJ, Proudfoot NJ. 2009. Pre-mRNA Processing Reaches Back toTranscription and Ahead to Translation. Cell. 136(4):688–700. doi:10.1016/j.cell.2009.02.001.

Morris JH, Apeltsin L, Newman AM, Baumbach J, Wittkop T, Su G, Bader GD, Ferrin TE. 2011. clusterMaker: a multi-algorithm clustering plugin for Cytoscape. BMC Bioinformatics. 12:436. doi:10.1186/1471-2105-12-436.

Nakamura A, Seydoux G. 2008. Less is more: specification of the germline by transcriptional repression. Development. 135(23):3817–3827. doi:10.1242/dev.022434.

Nishikura K. 2016. A-to-I editing of coding and non-coding RNAs by ADARs. Nat Rev Mol Cell Biol. 17(2):83–96. doi:10.1038/nrm.2015.4.

Payán-Bravo L, Fontalva S, Peñate X, Cases I, Guerrero-Martínez JA, Pareja-Sánchez Y, Odriozola-Gil Y, Lara E, Jimeno-González S, Suñé C, et al. 2021. Human prefoldin modulates co-transcriptional pre-mRNA splicing. Nucleic Acids Research. 49(11):6267–6280. doi:10.1093/nar/gkab446.

Pereira B, Billaud M, Almeida R. 2017. RNA-Binding Proteins in Cancer: Old Players and New Actors. Trends Cancer. 3(7):506–528. doi:10.1016/j.trecan.2017.05.003.

Perez-Perri JI, Noerenberg M, Kamel W, Lenz CE, Mohammed S, Hentze MW, Castello A. 2021. Global analysis of RNA-binding protein dynamics by comparative and enhanced RNA interactome capture. Nat Protoc. 16(1):27–60. doi:10.1038/s41596-020-00404-1.

Prashad S, Gopal PP. 2021. RNA-binding proteins in neurological development and disease. RNA Biology. 18(7):972–987. doi:10.1080/15476286.2020.1809186.

Ray D, Laverty KU, Jolma A, Nie K, Samson R, Pour SE, Tam CL, von Krosigk N, Nabeel-Shah S, Albu M, et al. 2023. RNA-binding proteins that lack canonical RNA-binding domains are rarely sequence-specific. Sci Rep. 13(1):5238. doi:10.1038/s41598-023-32245-9.

Richard S, Gross L, Fischer J, Bendalak K, Ziv T, Urim S, Choder M. 2021. Numerous Post-translational Modifications of RNA Polymerase II Subunit Rpb4/7 Link Transcription to Post-transcriptional Mechanisms. Cell Reports. 34(2):108578. doi:10.1016/j.celrep.2020.108578.

Ripin N, Parker R. 2023. Formation, function, and pathology of RNP granules. Cell. 186(22):4737–4756. doi:10.1016/j.cell.2023.09.006.

Roundtree IA, Evans ME, Pan T, He C. 2017. Dynamic RNA Modifications in Gene Expression Regulation. Cell. 169(7):1187–1200. doi:10.1016/j.cell.2017.05.045.

Shannon P, Markiel A, Ozier O, Baliga NS, Wang JT, Ramage D, Amin N, Schwikowski B, Ideker T. 2003. Cytoscape: A Software Environment for Integrated Models of Biomolecular Interaction Networks. Genome Res. 13(11):2498–2504. doi:10.1101/gr.1239303.

Singh S, Vanden Broeck A, Miller L, Chaker-Margot M, Klinge S. 2021. Nucleolar maturation of the human small subunit processome. Science. 373(6560):eabj5338. doi:10.1126/science.abj5338.

Smith E, Shilatifard A. 2010. The Chromatin Signaling Pathway: Diverse Mechanisms of Recruitment of Histone-Modifying Enzymes and Varied Biological Outcomes. Molecular Cell. 40(5):689–701. doi:10.1016/j.molcel.2010.11.031.

Smith RN, Aleksic J, Butano D, Carr A, Contrino S, Hu F, Lyne M, Lyne R, Kalderimis A, Rutherford K, et al. 2012. InterMine: a flexible data warehouse system for the integration and analysis of heterogeneous biological data. Bioinformatics. 28(23):3163–3165. doi:10.1093/bioinformatics/bts577.

Statello L, Guo C-J, Chen L-L, Huarte M. 2021. Gene regulation by long non-coding RNAs and its biological functions. Nat Rev Mol Cell Biol. 22(2):96–118. doi:10.1038/s41580-020-00315-9.

Sternberg PW, Van Auken K, Wang Q, Wright A, Yook K, Zarowiecki M, Arnaboldi V, Becerra A, Brown S, Cain S, et al. 2024. WormBase 2024: status and transitioning to Alliance infrastructure. Genetics. 227(1):iyae050. doi:10.1093/genetics/iyae050.

Szklarczyk D, Gable AL, Lyon D, Junge A, Wyder S, Huerta-Cepas J, Simonovic M, Doncheva NT, Morris JH, Bork P, et al. 2019. STRING v11: protein–protein association networks with increased coverage, supporting functional discovery in genome-wide experimental datasets. Nucleic Acids Research. 47(D1):D607–D613. doi:10.1093/nar/gky1131.

Szklarczyk D, Kirsch R, Koutrouli M, Nastou K, Mehryary F, Hachilif R, Gable AL, Fang T, Doncheva NT, Pyysalo S, et al. 2023. The STRING database in 2023: protein–protein association networks and functional enrichment analyses for any sequenced genome of interest. Nucleic Acids Research. 51(D1):D638–D646. doi:10.1093/nar/gkac1000.

Tafforeau L, Zorbas C, Langhendries J-L, Mullineux S-T, Stamatopoulou V, Mullier R, Wacheul L, Lafontaine DLJ. 2013. The Complexity of Human Ribosome Biogenesis Revealed by Systematic Nucleolar Screening of Pre-rRNA Processing Factors. Molecular Cell. 51(4):539–551. doi:10.1016/j.molcel.2013.08.011.

Tamburino AM, Ryder SP, Walhout AJM. 2013. A Compendium of Caenorhabditis elegans RNA Binding Proteins Predicts Extensive Regulation at Multiple Levels. G3 Genes|Genomes|Genetics. 3(2):297–304. doi:10.1534/g3.112.004390.

Tang H-W, Hu Y, Chen C-L, Xia B, Zirin J, Yuan M, Asara JM, Rabinow L, Perrimon N. 2018. The TORC1-Regulated CPA Complex Rewires an RNA Processing Network to Drive Autophagy and Metabolic Reprogramming. Cell Metabolism. 27(5):1040–1054.e8. doi:10.1016/j.cmet.2018.02.023.

Tapial J, Ha KCH, Sterne-Weiler T, Gohr A, Braunschweig U, Hermoso-Pulido A, Quesnel-Vallières M, Permanyer J, Sodaei R, Marquez Y, et al. 2017. An atlas of alternative splicing profiles and functional associations reveals new regulatory programs and genes that simultaneously express multiple major isoforms. Genome Res. 27(10):1759–1768. doi:10.1101/gr.220962.117.

Teigelkamp S, Achsel T, Mundt C, Göthel SF, Cronshagen U, Lane WS, Marahiel M, Lührmann R. 1998. The 20kD protein of human [U4/U6.U5] tri-snRNPs is a novel cyclophilin that forms a complex with the U4/U6-specific 60kD and 90kD proteins. RNA. 4(2):127–141.

Ule J, Blencowe BJ. 2019. Alternative Splicing Regulatory Networks: Functions, Mechanisms, and Evolution. Molecular Cell. 76(2):329–345. doi:10.1016/j.molcel.2019.09.017.

Utriainen M, Morris JH. 2023. clusterMaker2: a major update to clusterMaker, a multi-algorithm clustering app for Cytoscape. BMC Bioinformatics. 24(1):1–28. doi:10.1186/s12859-023-05225-z.

Vaishali, Dimitrova-Paternoga L, Haubrich K, Sun M, Ephrussi A, Hennig J. 2021. Validation and classification of RNA binding proteins identified by mRNA interactome capture. RNA. 27(10):1173–1185. doi:10.1261/rna.078700.121.

Velázquez-Cruz A, Baños-Jaime B, Díaz-Quintana A, De la Rosa MA, Díaz-Moreno I. 2021. Post-translational Control of RNA-Binding Proteins and Disease-Related Dysregulation. Front Mol Biosci. 8. doi:10.3389/fmolb.2021.658852. [accessed 2024 Sep 25]. https://www.frontiersin.org/journals/molecular-biosciences/articles/10.3389/fmolb.2021.658852/full.

Warner AD, Gevirtzman L, Hillier LW, Ewing B, Waterston RH. 2019. The C. elegans embryonic transcriptome with tissue, time, and alternative splicing resolution. Genome Res. 29(6):1036–1045. doi:10.1101/gr.243394.118.

Weinreb A, Varol E, Barrett A, McWhirter RM, Taylor SR, Courtney I, Basavaraju M, Poff A, Tipps JA, Collings B, et al. 2024. Alternative splicing across the C. elegans nervous system. :2024.05.16.594567. doi:10.1101/2024.05.16.594567. [accessed 2024 Oct 28]. https://www.biorxiv.org/content/10.1101/2024.05.16.594567v2.

Wilkinson ME, Charenton C, Nagai K. 2020. RNA Splicing by the Spliceosome. Annu Rev Biochem. 89:359–388. doi:10.1146/annurev-biochem-091719-064225.

Williams T, Ngo LH, Wickramasinghe VO. 2018. Nuclear export of RNA: Different sizes, shapes and functions. Seminars in Cell & Developmental Biology. 75:70–77. doi:10.1016/j.semcdb.2017.08.054.

Wolfe Z, Liska D, Norris A. 2024. Deep Transcriptomics Reveals Cell-Specific Isoforms of Pan-Neuronal Genes. :2024.05.16.594572. doi:10.1101/2024.05.16.594572. [accessed 2024 Oct 28]. https://www.biorxiv.org/content/10.1101/2024.05.16.594572v1.

Wolozin B, Ivanov P. 2019. Stress granules and neurodegeneration. Nat Rev Neurosci. 20(11):649–666. doi:10.1038/s41583-019-0222-5.

Zhao Y, Mir C, Garcia-Mayea Y, Paciucci R, Kondoh H, LLeonart ME. 2022. RNA-binding proteins: Underestimated contributors in tumorigenesis. Seminars in Cancer Biology. 86:431–444. doi:10.1016/j.semcancer.2022.01.010.

