## Supplementary material for "A global view of the RNA-binding and regulatory protein landscape in *Caenorhabditis elegans*": Figure S1

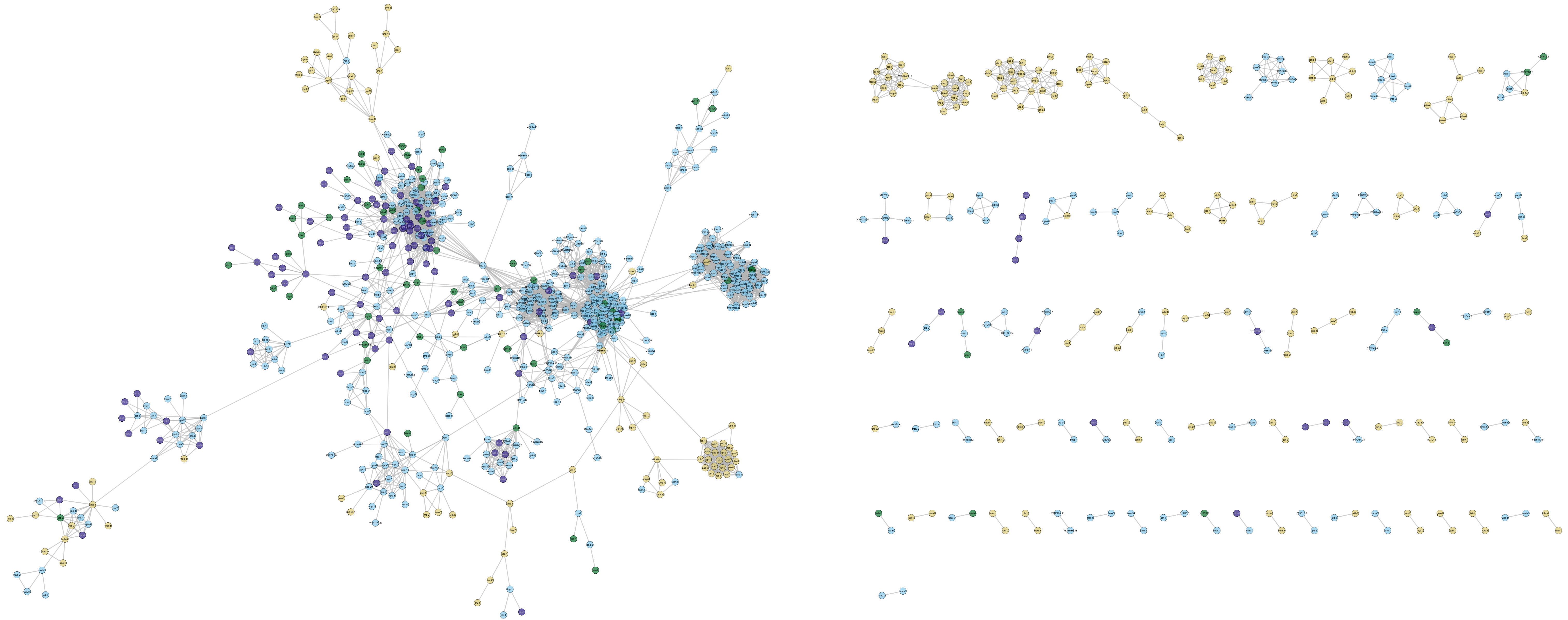

**Figure S1.** RNA regulatory protein-protein interaction network comprising Class 1-4 proteins, with nodes labelled with gene names. Related to Figure 2.
