## Supplementary material for "A global view of the RNA-binding and regulatory protein landscape in *Caenorhabditis elegans*": Figure S2

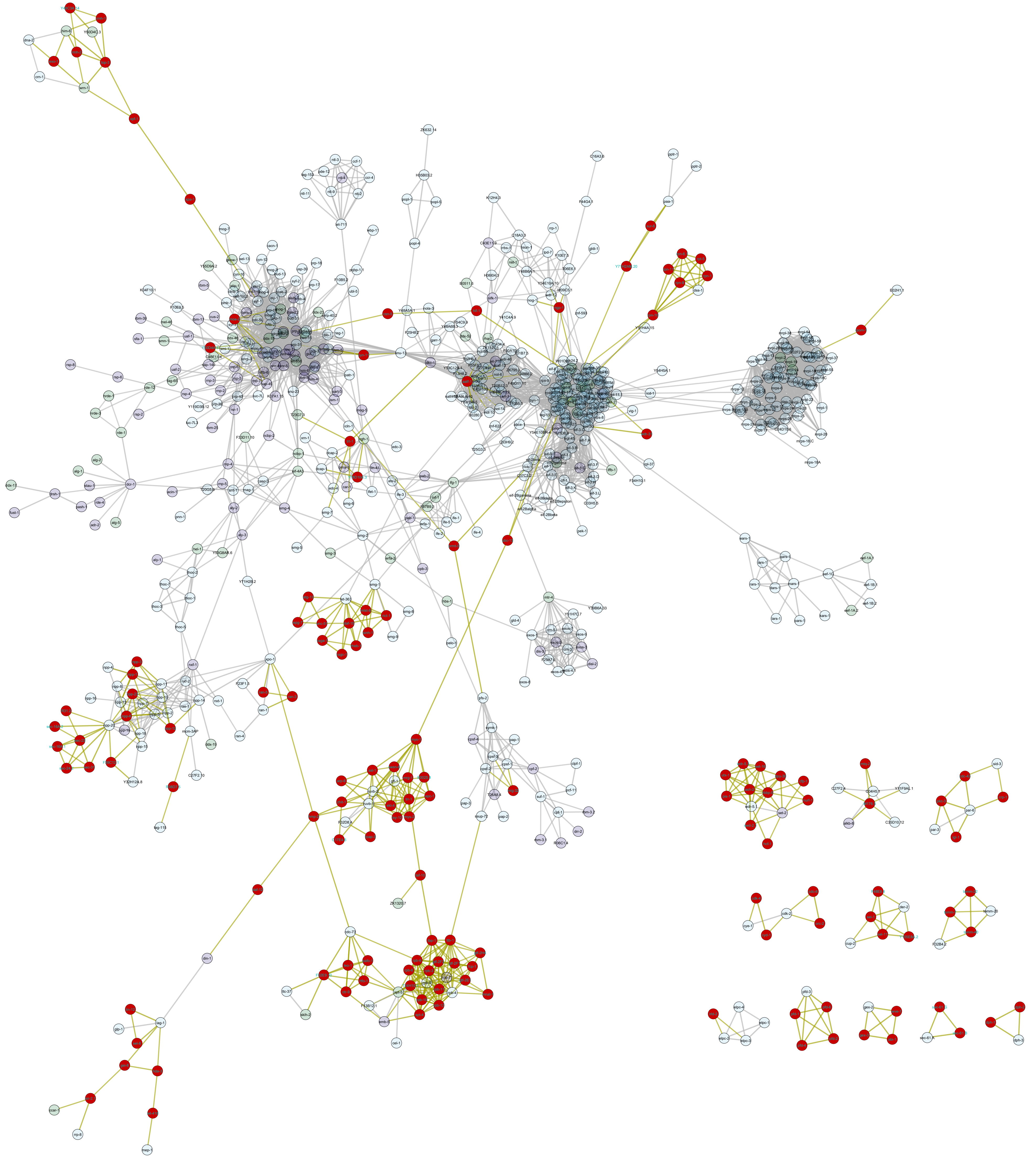

**Figure S2.** Regulatory protein-protein interaction network comprising Class 1-3 and Class 5 proteins, with nodes labelled with gene names. Related to Figure 3.
