## Supplementary material for "A global view of the RNA-binding and regulatory protein landscape in *Caenorhabditis elegans*": Figure S3

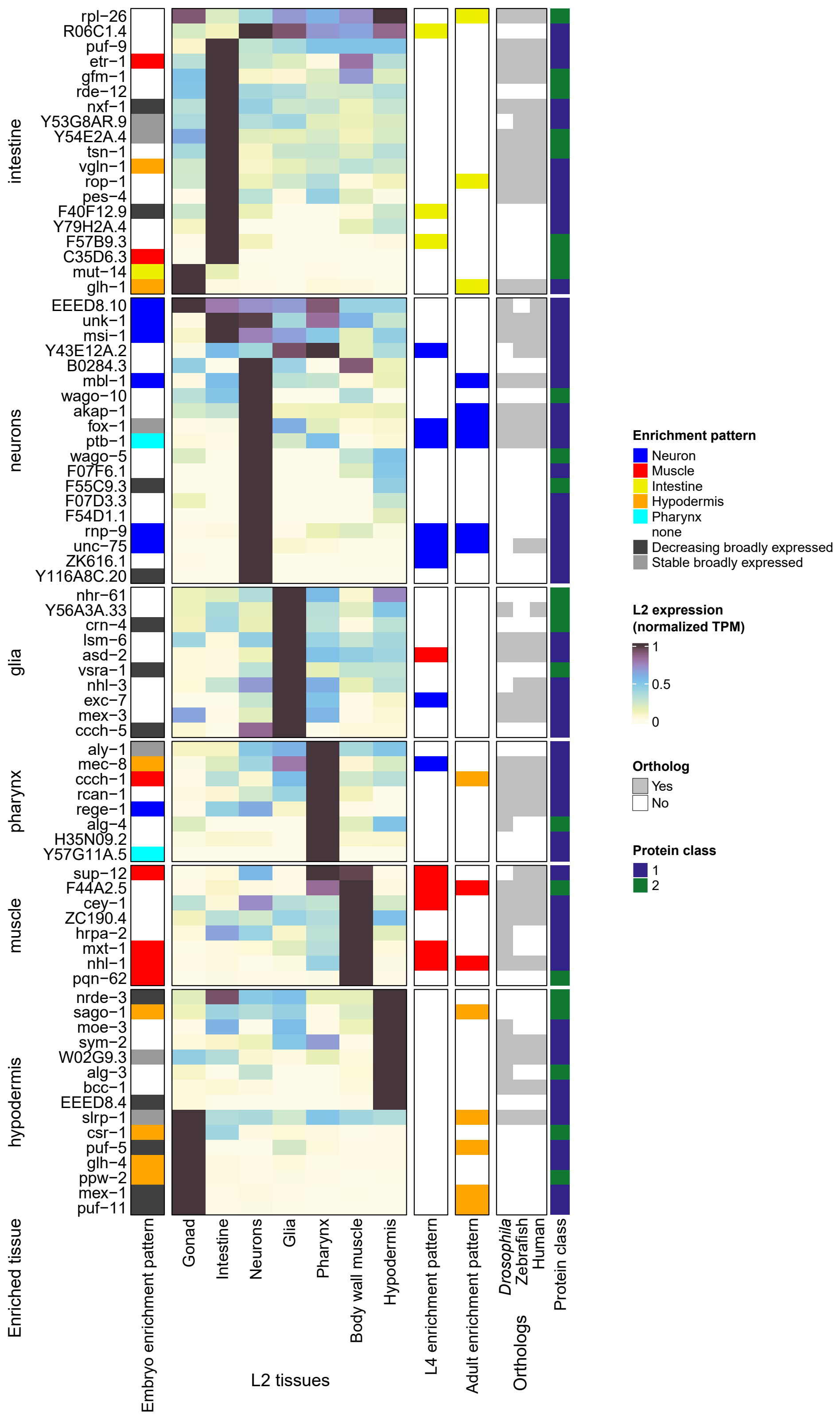


**Figure S3. Overview of highly tissue-enriched RNA regulatory proteins in classes 1 and 2.** Summary of expression of Class 1 and 2 RNA regulatory proteins that are enriched for expression in a particular somatic tissue at one or more developmental stages. Data shown are as in Figure 4A: tissue-specific expression data in L2 larvae from Cao et al. 2017; descriptions of tissue enrichment or expression patterns in the embryonic, L4 larval, and adult stages, from Warner et al. 2019, Gracida et al. 2017, and Kaletsky et al. 2018, respectively; the presence or absence of a direct ortholog of each *C. elegans* protein in *Drosophila melanogaster*, zebrafish, or humans; and the corresponding RNA regulatory protein class. Proteins were grouped according to primary tissue of expression based first on expression patterns in L2 larvae, then enriched tissue annotations in L4 larvae, adults, and embryos, respectively.
